# Resource-explicit interactions in spatial population models

**DOI:** 10.1101/2024.01.13.575512

**Authors:** Samuel E. Champer, Bryan Chae, Benjamin C. Haller, Jackson Champer, Philipp W. Messer

## Abstract

Continuous-space population models can yield significantly different results from their panmictic counterparts when assessing evolutionary, ecological, or population-genetic processes. However, the computational burden of spatial models is typically much greater than that of panmictic models due to the overhead of determining which individuals interact with one another and how strongly they interact. Though these calculations are necessary to model local competition that regulates the population density, they can lead to prohibitively long runtimes. Here, we present a novel modeling method in which the resources available to a population are abstractly represented as an additional layer of the simulation. Instead of interacting directly with one another, individuals interact indirectly via this resource layer. We find that this method closely matches other spatial models, yet can dramatically increase the speed of the model, allowing the simulation of much larger populations. Additionally, models structured in this manner exhibit other desirable characteristics, including more realistic spatial dynamics near the edge of the simulated area, and an efficient route for modeling more complex heterogeneous landscapes.

## INTRODUCTION

A common abstraction when modeling evolutionary processes is to treat populations as well-mixed and randomly mating. This assumption of so-called “panmixia” often allows for tractable mathematical solutions and efficient simulation approaches, and therefore lies at the core of many population models^1,2^. Yet, while panmixia may be a suitable assumption for describing populations in laboratory settings, small islands, and a handful of other cases^3–5^, there are many contexts in the natural world that can only be accurately investigated with some representation of spatiality^6–9^.

One approach for incorporating spatial population structure is to break up a population into discrete subpopulations that are treated as separate panmictic demes linked by migration^7,10–12^. This can be a good approximation when modeling separate islands with inter-island dispersal, landscapes with discrete habitat patches, and similar scenarios. However, it may not be well-suited to populations that inhabit larger continuous landscapes. For such populations, it is often critical to model continuous space explicitly to capture key aspects of the evolutionary dynamics, such as the wave of advance of a strongly beneficial allele or the spread of an invasive species where it has been newly introduced^8,13–17^.

Unfortunately, individual-based simulations of continuous-space populations are typically marked by much longer runtimes than panmictic models, often rendering them a less desirable or outright impractical choice. These longer runtimes are a result of the significant computational cost of spatial calculations. Throughout the simulation, spatial positions are used to make determinations such as which individuals can mate with one another and which are in competition, and these determinations must be updated whenever individuals move, die, or are born. To make these determinations, each individual must ascertain whether or not it is interacting with each other individual; if two individuals interact, the strength of the interaction, usually a function of the distance between the individuals, must then be computed (we will call these “direct-interaction models”). Superficially, it might appear that this process would require on the order of *n*^2^ operations for a population of *n* individuals, but since individuals are usually defined as only interacting when within some threshold distance, the use of a space-partitioning data structure to search for nearby individuals can reduce the workload substantially. The more spatially local the interactions, the greater the performance improvement yielded by such data structures^18,19^. Nonetheless, spatial calculations still often occupy the majority of the runtime of direct-interaction models, and can thus be the main factor that prevents the use of continuous-space models or limits their practical scale.

Additional problems crop up near the boundaries of spatial models. If individuals are distributed approximately uniformly, as might be considered appropriate in many cases, individuals at the edges of the simulated area tend to interact with fewer neighbors. Since spatial population regulation depends on local density, this can result in edge areas becoming overpopulated or other deviations from desired model behavior. Such edge effects can in principle be corrected by calculating local density only within the modeled area^20^. However, this correction is not always trivial, and thus represents an additional computational burden, especially when interaction strength is governed by more complex functions of distance, or when modeling an area with an irregular border (such as an island’s coastline) rather than a square or rectangular area.

In this study, we introduce a new paradigm for individual-based simulations on continuous-space landscapes in which competitive interactions can be evaluated much faster, and which does not require a computational correction for edge effects. This is accomplished by explicitly simulating the resources available to each individual. These resources are abstractly modeled as “resource nodes” which are distributed across the landscape either uniformly or randomly. Competing individuals collect resources from the nodes in their foraging area, reducing the resources available to other individuals. In this way, the local density of individuals is regulated indirectly, through extrinsic resource availability, rather than by direct interactions between individuals. With an appropriately chosen resource-node density, each resource node can support numerous individuals.

Since individuals in this model interact with resource nodes instead of with one another, and given that there are fewer nodes than individuals, competition is determined using fewer total spatial interactions as compared to direct-interaction models, potentially resulting in a substantially faster simulation. Model dynamics near edge areas of resource-explicit models may also be preferable to those in direct-interaction models. Individuals at the edges of the landscape have fewer local competitors, but they are also near proportionately fewer resource nodes (since there are neither competitors nor nodes outside of the modeled area). Consequently, local carrying-capacity density is the same near the edges of the area as it is in the interior of the landscape without the need for any time-consuming corrections, even when the edge of the landscape has a complex, natural shape like a coastline.

Our resource-explicit method results in interaction strengths that qualitatively differ from directly calculated individual–individual interaction strengths. Most notably, interaction strengths are not a rigidly defined function of distance, but can rather be described as having a distribution of possible values at a given distance. This characteristic might be considered biologically realistic, since many competitive interactions in real biological systems are mediated by an extrinsic resource, and the intensity with which a given pair of individuals compete is likely influenced by many factors and can vary between different pairs of individuals even when they are separated by the same distance. We show that the variance of the distribution of interaction strengths can be narrowed to a desired range by selecting a sufficiently high resource-node density (at some cost to performance).

Furthermore, we also show that models using this technique can run faster (in some of our simulation scenarios over 20 times faster) than direct-interaction models when modeling a dense population (approximately matching the density of an urban population of rats^21^). Given this speedup, our resource-mediated interaction approach can allow simulations to be scaled up well beyond the practical limits of direct-interaction models, opening up new possibilities for biological realism.

Finally, we will argue that resource-explicit models offer an elegant implementation path for a number of desirable features, including heterogeneous landscapes, irregular spatial boundaries such as coastlines, and efficiently calculated interactions among multiple species.

## METHODS

In order to facilitate a direct comparison between the resource-explicit technique introduced here and commonly used direct-interaction models, a simple population model was developed to serve as a shared platform. This core model, and the derivations from it described below, are written in the SLiM individual-based forward-time evolutionary modeling framework (version 4.1)^20^.

Our core model implements a population in which competition determines mortality. Individuals inhabit a two-dimensional continuous-space landscape, consisting of a square area with reprising boundaries (i.e., dispersal coordinates are re-drawn when those coordinates fall outside the defined area). The simulated area is measured in multiples of a basic “unit area,” defined as the foraging area of an individual of the species being modeled. The foraging radius, *r*, is thus defined as 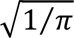 (such that the area foraged equals 1), and the maximum distance at which two individuals can interact is defined as 2*r*, since this is the maximum distance at which there is any overlap between the foraging areas of two individuals. The overall size of the modeled area is controlled by a landscape-size parameter. The carrying-capacity density (carrying capacity per unit area) is controlled by a density parameter.

In each time-step, or “tick”, of the model, individuals first reproduce, and then experience competition-dependent mortality. Even in the resource-explicit models (with the exception of one variant described below), mate search is conducted using a direct interaction between males and females (with a maximum interaction distance of 2*r*) rather than an indirect resource-explicit interaction, since mate-search interactions are generally less of a runtime burden than competitive interactions because of the sex-constrained nature of the mate-search interaction. A fixed-strength interaction function is used for this purpose, resulting in each female having an equal probability to choose any male within range. Each female that reproduces generates a number of offspring drawn from a Poisson distribution with a mean of eight (arbitrarily chosen, but unimportant for our purposes here). Newly generated offspring are assigned a spatial position by starting at the same *x* and *y* coordinate as their maternal parent and then deviating on each axis by a value drawn from a normal distribution with a mean of zero and a standard deviation of *2r*.

Starting from this shared model platform, we derived several model variations differing only in their implementation of competition. Three of the variants are regulated by direct individual–individual interactions using three frequently used interaction functions. These three direct-interaction models are used as a baseline against which we compared four resource-explicit model variants. Finally, we also prepared a panmictic model in order to assess the proportion of runtime spent by the other models on performing spatial computations (as opposed to other necessary functions such as offspring generation).

### I. Models <u>with</u> Direct Interactions between Individuals

There are several functions that are commonly used to determine the interaction strength between individuals in spatial models (Figure 1, left panel). The simplest is a fixed-strength interaction function: all individuals within range interact with the maximum possible strength (which we will call a strength of 1). Individuals at the maximum interaction range of 2*r* thus compete just as intensely as individuals that are closer to one another. This is usually an undesirable choice when simulating competition for resources because it can lead to clustering artifacts that may be unrealistic^22^. However, this function is the fastest to evaluate. A second option is a linear function: two individuals in the exact same location interact at strength 1, and that interaction strength linearly declines to 0 at the maximum interaction distance. Another option is a Gaussian function, in which two individuals in the same location interact at strength 1, with interaction strength declining non-linearly with distance, implemented in our model as:

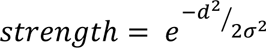

**Figure 1.**
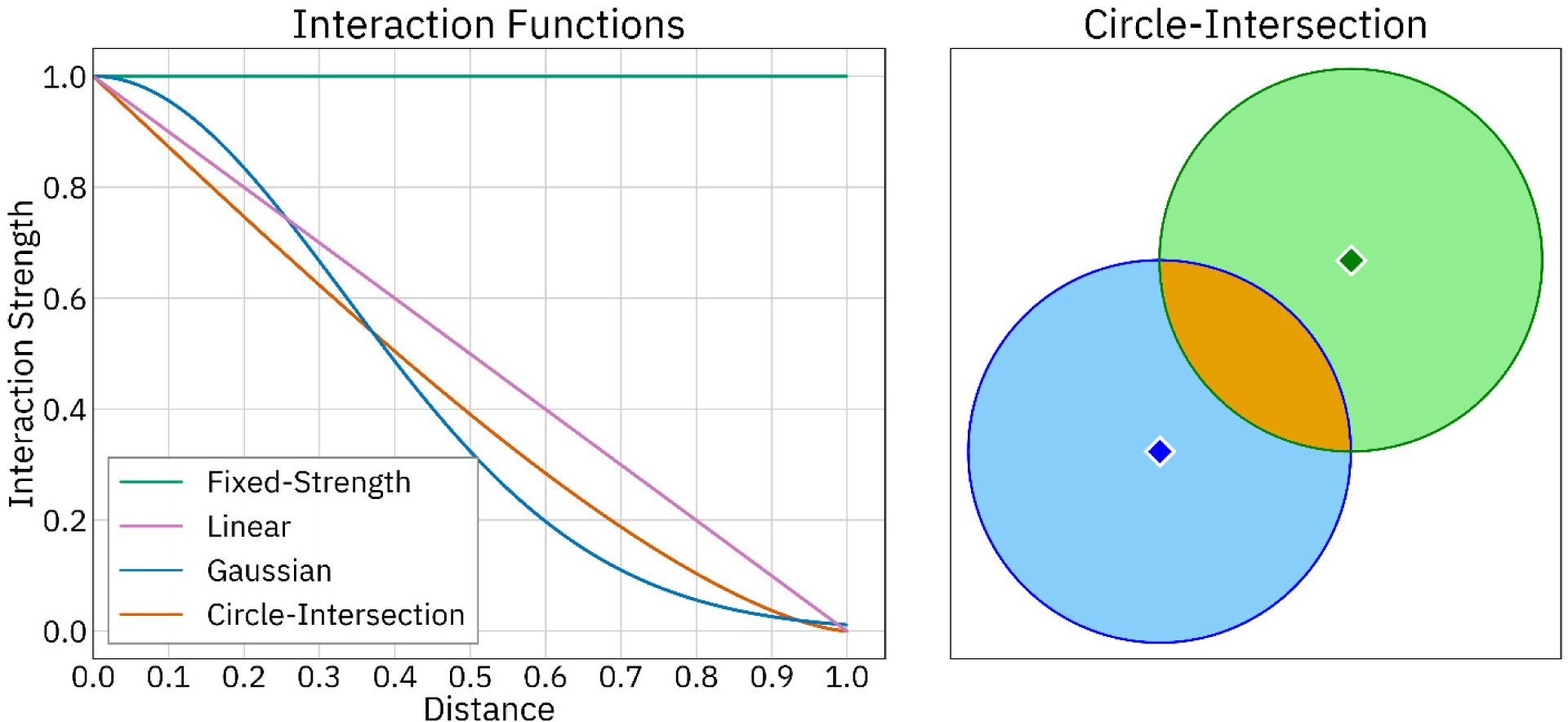
Interaction functions. Individuals in continuous-space models typically interact with one another according to an interaction strength function. Several interaction functions are depicted in the left-hand panel. The right-hand panel depicts competition between two individuals (blue and green diamonds with white outlines) using the circle-intersection function; the circle-intersection function was chosen for illustration because it is the most similar to the interaction strengths produced by our resource-explicit models. The foraging areas of the individuals are represented by blue and green shading; competition strength is proportional to the amount by which these foraging areas overlap (orange shaded area).

Where *d* is the distance between the two interactors as a fraction of the maximum interaction distance, and σ is the standard deviation of the interaction function (commonly called its “width”), which determines how quickly interaction strength decays as distance increases.

Each of these three interaction functions is incorporated into a separate model in this manuscript. For the Gaussian interaction, we used a σ value of 1/3 of the maximum interaction distance to ensure that the interaction strength declines to a suitably small value (0.011) at the maximum interaction distance. After calculating competition between individuals, mortality for each individual is implemented as the sum of all competition experienced divided by the amount expected if the density of individuals in the local area were at carrying capacity.

In addition to these three common choices, many other functions can be used to calculate interaction strength. One function that is relevant here, although infrequently used, is the “circle-intersection” function. Given that all of the interaction functions considered above define each individual as foraging from a circular area, this function defines the interaction strength between two individuals to be equal to the intersection of their foraging areas (Figure 1, right panel). The resulting function has a strength of 1 for individuals that are in the exact same location, decreasing to 0 at the maximum interaction distance of 2*r* (at which range the foraging areas of the two individuals intersect at a single point), as described by:

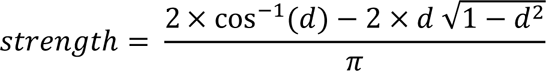

where *d* is the distance between individuals as a fraction of the maximum interaction distance. This function is not commonly used in spatial population models (and was not implemented in any of the models used in this manuscript), perhaps due to the amount of computation required. However, it is a well-behaved continuous function based on biologically plausible assumptions, and is mentioned here because it provides a useful point of comparison with the resource-explicit models presented next.

We also include a panmictic model in which individuals are not assigned spatial coordinates. Instead of local competition, a global survival rate is imposed on the basis that all individuals equally compete with all other individuals. This survival rate is calculated as the carrying capacity of the system divided by the current population size. Instead of choosing a nearby mate, females randomly select a male from the population as a whole. The panmictic model is otherwise identical to the other models.

### II. Resource-Explicit Models

In the resource-explicit models, the landscape is populated with resource nodes at the outset of the simulation. The number of nodes to be placed on the landscape depends on the desired node density and the size of the landscape. For example, a landscape with a total area of 1,000 times the unit area (the foraging area of an individual) populated with a node density of 25 nodes per unit area will be populated with 25,000 nodes.

Interactions between individuals in the resource-explicit models could be considered analogous to interactions calculated by the circle-intersection function, since individuals in these models forage from nearby nodes in a circular area, and competition between two individuals is proportional to how much their ranges overlap (in terms of how many nodes are shared between them). Two individuals in the same location forage from the same resource nodes, and could thus be considered to have an interaction strength of 1. That interaction strength decreases as the distance between the individuals increases, because the number of nodes they share decreases.

Resource nodes are assigned a value representing the amount of resources available in the local area, as determined by the density parameter of the modeled species as well as the density of the resource nodes. To continue the previous example, if the population has a capacity density of 100 individuals per unit area and nodes have a density of 25 per unit area, then each node has resource value of 4.

The resource-explicit models developed in this study were each tested with three different resource-node placement methods: a uniform hexagonal tiling of the landscape, a uniform square tiling of the landscape, and a random placement of nodes across the landscape, with positions re-randomized each tick of the model. Each of these placement methods was tested with a density of 12, 25, and 50 nodes per unit area. Although resource nodes may conceptually consist of a two-dimensional area (and are depicted as polygons in some figures), the resource nodes in the model are point entities.

In the SLiM models presented in this manuscript, resource nodes are implemented as a second species for which no life cycle is defined (i.e., no mortality or reproduction) that spatially interacts with the modeled individuals. Similar implementations may be possible in other modeling frameworks.

See “Resource Node Placement Methods” in the supplement for a more detailed description of these methods, along with guidance on choosing node density and placement method.

#### II. A. Competition in Resource-Explicit Models

We implemented two alternative methods governing how foraging competition is calculated within the resource-explicit framework. In the first of these, individuals forage from an “inelastic” foraging area, only gathering resources from within a fixed distance. In the second method, individuals forage from an “elastic” range, and can potentially forage from further away in order to maintain a full-sized foraging area.

In the “inelastic” method (Figure 2, left panels; see Supplemental Figures 1 and 2 for versions of this figure that use a hexagonal tiling and a random distribution of resource nodes, respectively), individuals forage from all of the resource nodes that are within their foraging radius, *r*. Competition in the inelastic model is calculated by first iterating through each resource node. Each node tallies local demand as the number of individuals within distance *r* and then sends each such individual an amount of resources equal to the total amount at that node divided by the local demand. After each node has distributed its resources, individuals are assigned a probability of survival according to their amount of resources received, as described in the following example.

**Figure 2.**
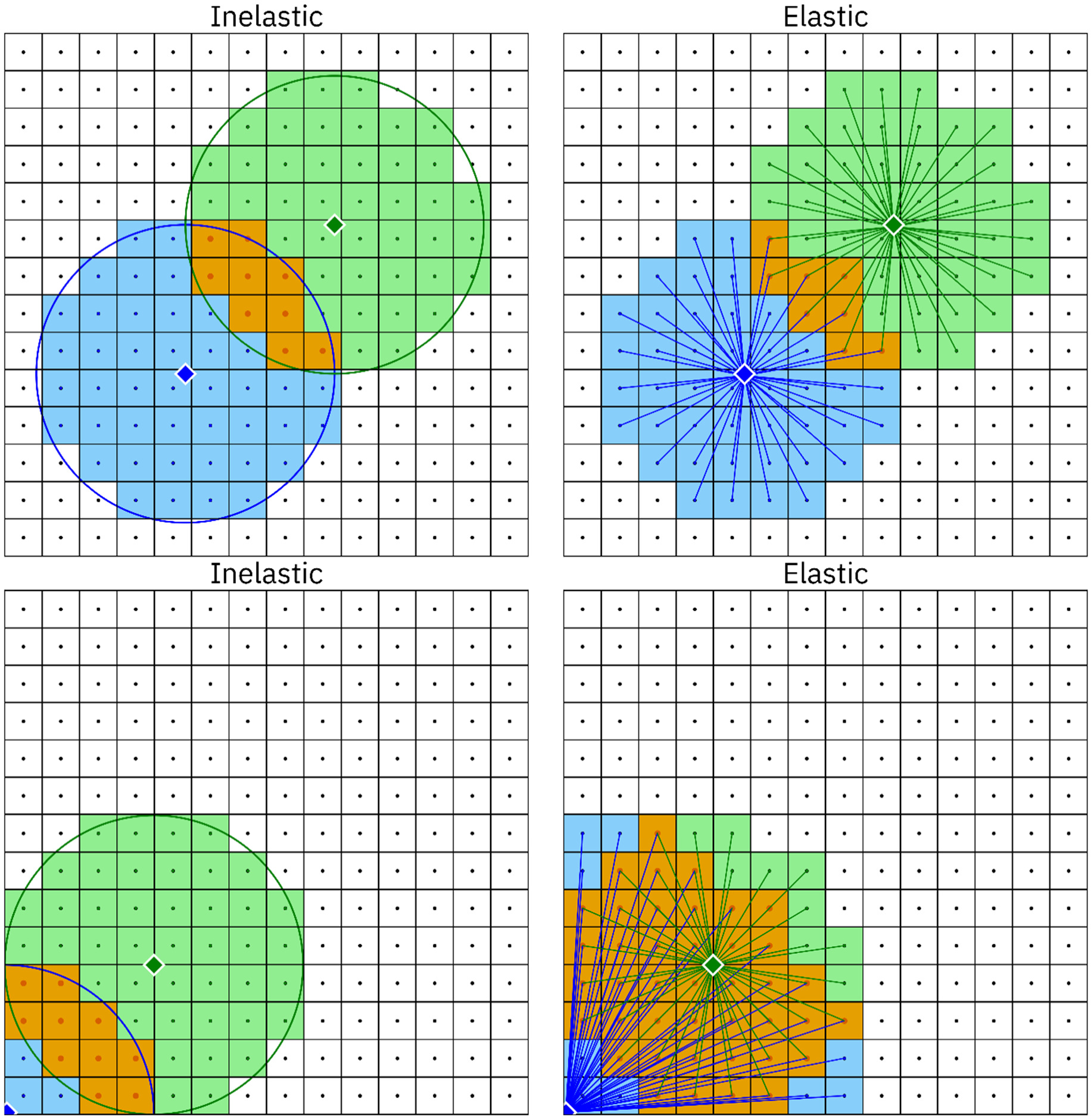
Visualization of resource-explicit interaction algorithms, square tiling. Competition between two individuals (blue and green diamonds) is determined by the portion of their foraging area that overlaps and by the interaction algorithm used in the model. The foraging areas are represented by blue and green shading; the overlapping area is shaded orange. In the inelastic model (left), individuals forage from resource nodes (dots at the center of each square) within their foraging radius. In the elastic model (right), individuals forage from as many nodes as necessary to maintain a nominally sized foraging area (in this case comprising 50 nodes). Away from the edges of the landscape (top row), the two models behave similarly but not identically: in the inelastic model (upper left panel), the blue individual happens to forage from 51 nodes (due to its precise spatial position within the grid), resulting in slightly greater competition. The difference between the two models is much greater in the corner of the landscape (bottom row): the blue individual has a much smaller area in the inelastic model, whereas the blue individual in the elastic model forages from much further away to maintain a full-sized foraging area, resulting in greater competition.

If the resource node density is 25 nodes per unit area and the carrying-capacity density of the modeled species is 100 individuals per unit area, then each node has a total resource value of 4. If the current population size is five times the total carrying capacity of the system, and individuals are uniformly distributed, then each of the 25 nodes an individual forages from provides an average of 4 / 500 resources (supply at the node divided by the average demand). After collecting resources from 25 such nodes, individuals will have collected an average resource amount of (4 / 500) *×* 25 = 1 / 5. The amount of resources collected is directly used as a survival rate (20%, in this case), which will, on average, bring the population to its carrying capacity.

In models using the inelastic method, individuals are not guaranteed to have access to the full number of resource nodes that corresponds to the defined foraging area (the “nominal” foraging area of the species). Indeed, individuals in the corners of the landscape might forage from only a quarter of their nominal foraging area (Figure 2, bottom left panel; see Supplemental Figures 1 and 2). Even individuals in the interior of the landscape may have access to somewhat more or fewer nodes (for example, the blue individual in the upper left panel of Figure 2 forages from 51 nodes, while the nominal foraging area is 50 nodes). Individuals in the interior of the landscape with fewer nodes in range have proportionately lower rates of survival, whereas individuals with more nodes in range have proportionately higher rates of survival. Near the edge of the landscape, individuals have access to fewer nodes, but those nodes are also foraged at by fewer competitors, so these effects will tend to balance out.

In the “elastic” method (Figure 2, right panels; see Supplemental Figures 1 and 2), individuals forage from as many nodes as necessary in order to maintain their nominal foraging area – though they are limited to the nodes that exist within radius *R*, where *R* is greater than the foraging radius *r* in the inelastic model. Given a node density of *n* nodes per unit area, individuals will attempt to forage from the nearest *n* nodes. With a sufficiently large *R*, individuals will always maintain their full nominal foraging area (an *R* of two times *r* is sufficient given a square landscape, and this value was used in the models in this manuscript). In this way, individuals near the edges of the landscape are able to forage from further away to compensate for the lack of nodes in their vicinity, resulting in an elastic foraging area (Figure 2, bottom right panel; see Supplemental Figures 1 and 2).

In the elastic method, competition is determined as follows. First, all individuals are iterated through, and the nearest *n* resource nodes to each individual that are within radius *R* are recorded. In the process, every time a resource node is recorded, a variable belonging to the node that tracks local demand is incremented by one. After this, all individuals are iterated through once again. During this second iteration, each individual receives resources from each of their nodes according to the amount at the node divided by the local demand at that node. After receiving resources, individuals are assigned a survival probability according to the amount received, in the same manner described above for the inelastic method.

#### II.B. Additional Resource-Explicit Model Variants

In addition to the basic inelastic and elastic models above, we include two variants of the inelastic model. The changes that define these variants are compatible with one another, and could be combined in a single model if desired.

##### Fair variant

This variant corrects for the spatial variation in mortality rate found in the default inelastic model, such that the survival rate of an individual is unaffected by having access to more or fewer resource nodes. This is achieved by performing a preliminary iteration through the nodes and incrementing a variable for each individual that tracks the number of nodes it forages from (with this variable denoted *f_x_* for a given individual *x*). Then, the amount of resources an individual *i* receives from a given resource node is determined as follows:

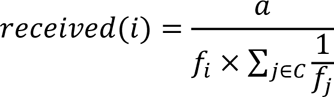

where *a* is the amount of resources available at the node, and *C* is the set of all competitors receiving resources from the node.

In this variant, conceptually, all individuals have an equal amount of time or energy to forage within their foraging area. Individuals with fewer nodes in range spend more time at each of those nodes, and thus gather more resources from them, while individuals with more nodes in range spend less time at each node.

##### Resource-explicit reproduction variant

This variant replaces the mate-search interaction with a resource-explicit interaction, thus replacing all direct interactions between individuals with interactions between individuals and resource nodes. Before reproduction takes place, each node caches a list of males that are within foraging range. Then, rather than directly searching for and choosing a nearby male, females instead select a mate from the concatenated lists of males cached at all of the nodes within foraging range.

The result of this approach is that potential mates are not chosen with equal probability, as in the other models. For example, if a male and female have ranges that only overlap at a single resource node, that male will be in the female’s list of candidate mates only a single time. A closer male who shares 25 nodes with the female will be duplicated in the list 25 times, resulting in a 25 times higher chance for the female to select the closer male. Note that similar dynamics can be implemented using a direct interaction with an appropriate interaction function.

## RESULTS

### I. Dynamics of the Resource-Explicit Models

The resource-explicit models replace direct competitive interactions between individuals with indirect competition for resources at discrete nodes located across the landscape. The resulting indirect interaction function differs from commonly used direct interaction functions in two qualitative ways. First, the indirect interaction function is discontinuous: as the distance between the two individuals increases, the number of nodes they share, and thus their interaction strength, decreases in discrete steps, not continuously. Second, the indirect interaction function is not position-invariant: two individuals at a given distance may interact with a different strength depending on their specific positions on the landscape with respect to the resource nodes.

However, despite these qualitative differences, a quantitative assessment reveals that interaction strengths in the resource-explicit models closely match those calculated using the circle-intersection interaction function. To compare interaction strengths between methods, a pair of individuals was placed at random locations on a large landscape, and interaction strengths between the individuals were measured using the circle-intersection function, the inelastic resource-explicit method, and the elastic resource-explicit method. This was repeated 2 million times in each of nine separate resource-node placement scenarios (hexagonal tiling, square tiling, and random placement, each at a density of 12, 25, and 50 nodes per unit area). Individuals were not placed near the edges of the landscape in order to avoid distortion produced by different edge dynamics. Each interaction strength measured using the circle-intersection function was then subtracted from the corresponding measurement from each resource-explicit model to yield distributions of differences (Figure 3, Supplemental Figures 4 and 5; see Supplemental Figures 8-13 for visualizations of the resource-explicit functions).

**Figure 3.**
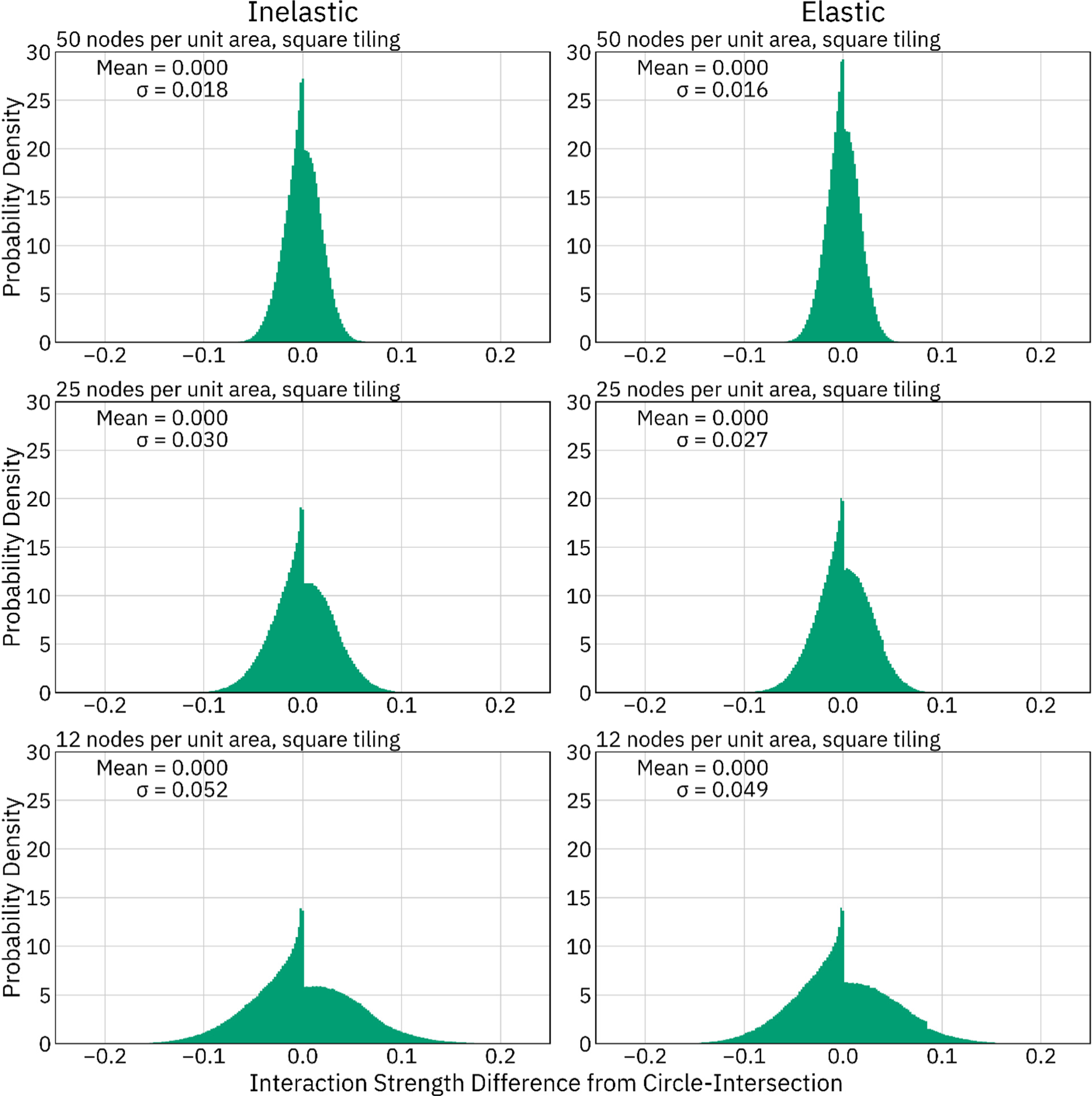
Differences in interaction strength between resource-explicit models and the circle-intersection function, square-tiled. Two million pairwise interaction strengths between randomly placed individuals as measured by the circle-intersection function were subtracted from those measured in resource-explicit models to yield a distribution of the deviation from the circle-intersection function for each model. As node density increases, the standard deviation decreases. These distributions all have a distinctive peak just below 0 due to cases where pairs of individuals have a very small but non-zero interaction strength when using the circle-intersection function, but the small overlapping portion of their foraging areas does not include any resource nodes.

This analysis found that the variation in interaction strength in the resource-explicit models decreases as the density of resource nodes increases. Furthermore, variance in the elastic model was lower than in the inelastic model in some cases. Using either a square or hexagonal tiling (Figure 3 and Supplemental Figure 4) the interaction strengths closely mirrored those calculated by the circle-intersection function, with a standard deviation in interaction strengths of 0.018 or less at a density of 50 nodes per unit area, and as high as 0.056 at a density of 12 nodes per unit area. When resource nodes were randomly placed (Supplemental Figure 5), the standard deviation in interaction strengths was substantially higher, measuring 0.049 and 0.054 at a density of 50 nodes per unit area in the elastic and inelastic models respectively, and 0.100 to 0.113 at a density of 12 nodes per unit area.

These distributions show that any given pairwise interaction strength in a resource-explicit model tends to be fairly close to the strength as calculated by the circle-intersection function. However, each individual in a spatial model typically interacts with many other individuals. These stochastic differences further average out over all of the interactions that determine the overall level of competition experienced by each individual. To quantify potential differences in overall competition experienced by individuals in the resource-explicit models, we conducted an additional analysis in which a population of 400,000 was simulated using the inelastic, “fair” inelastic, and elastic method at a density of 20 individuals per unit area. After the offspring generation step of the tenth tick of the simulation, the survival rates of the centermost 100,000 individuals were recorded (as before, only central individuals were selected in order to avoid distortion produced by different edge dynamics). The survival rates for each individual were then recalculated as would be done in a direct-interaction model using the circle-intersection function. The survival rates for each individual calculated by the circle-intersection function were then subtracted from those calculated using the resource-explicit function to yield a distribution of differences in survival rates experienced by the individuals in the model. This distribution of differences in survival rate reveals that the resource-explicit models match the circle-intersection interaction function very closely (Figure 4 and Supplemental Figures 6 and 7) – even more closely than the analysis that assessed pairwise interactions one at a time (Figure 3) indicates. At a density of 12 nodes per unit area, the standard deviation of the inelastic model was 0.018 using square tiling, 0.008 using hexagonal tiling, and 0.058 using randomly placed nodes. However, the standard deviation of the “fair” inelastic and the elastic model were much lower – no more than 0.002 in the regularly tiled models, though up to 0.034 in the model with randomly placed nodes.

**Figure 4.**
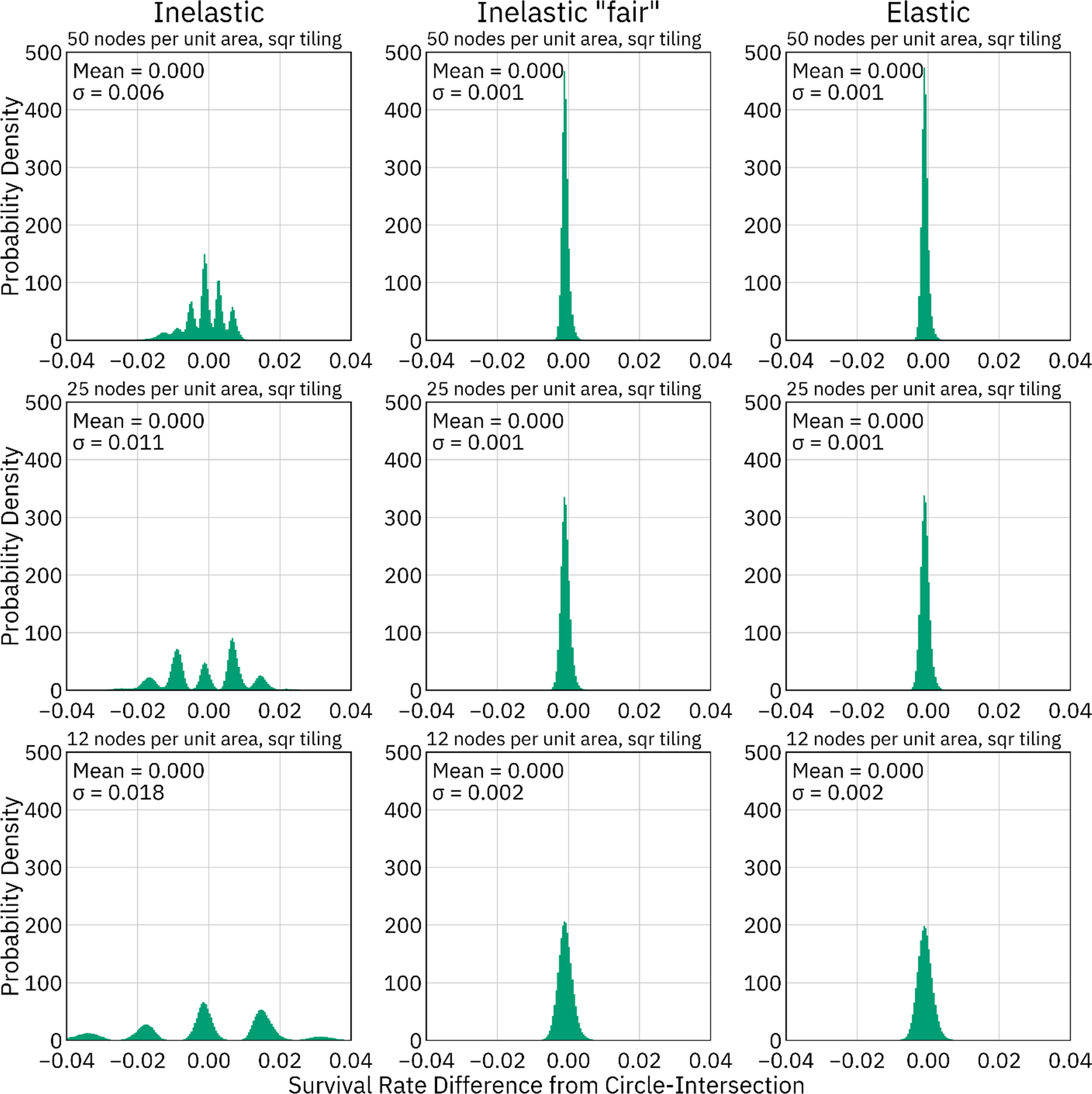
Differences in survival rate between resource-explicit models and a direct-interaction model using the circle-intersection function, square-tiled. The survival rates of 100,000 individuals were measured using the inelastic method, the “fair” inelastic method, the elastic method, and the circle-intersection function. Strengths measured by the circle-intersection function were subtracted from measurements made using the other resource-explicit methods for each individual to yield distributions of differences.

In the inelastic model, individuals sometimes forage from slightly more or fewer nodes than their nominal foraging area, resulting in a reduced or increased mortality rate. This is the reason for the multimodal distribution in the inelastic models in Figure 4 (and Supplemental Figures 6 and 7). For example, the central peak in the lower left panel is the distribution of survival rates for individuals who foraged from 12 nodes, while the peaks to the left and right of the central peak are the distributions of survival rates for individuals who foraged from 11 and 13 nodes, respectively. In the uniformly tiled models, this phenomenon could be described as a repeating, micro-scale habitat-quality gradient within each tile of the model (Supplemental Figures 14-19). In most cases, this variation in mortality rate can be disregarded as a negligible additional source of stochasticity compared to the stochasticity already involved in mortality determinations. However, this phenomenon may be more of an issue in sparse populations with small litter sizes. In such models, outcomes could be affected if a few individuals randomly experience premature mortality just because they happen to disperse into a particularly inhospitable micro-habitat within a tile. For modeling such systems, the “fair” variant which compensates for these effects (see methods) may be preferable because survival rates in this variant are almost identical to those in the elastic model (Figure 4 and Supplemental Figures 6 and 7).

### II. Computational Performance

To assess the runtime performance of each method, a series of simulations was conducted. For each model, 40 measurements of the elapsed runtime per tick were taken with each of two population sizes (100,000 and 1 million individuals), at each of two levels of population density (20 and 200 individuals per unit area), and at each of three levels of resource-node density (12, 25, and 50 nodes per unit area; Table 1). Simulations were performed on a desktop computer using an Intel i9-9900K CPU, on which only a single simulation was performed at a time in order to keep conditions across simulations as consistent as possible. Absolute runtimes in other contexts will vary. Memory usage of the models is not presented in detail here, but it was observed that the elastic models used up to three times as much memory as the direct-interaction models, while the default inelastic model, the “fair” variant, and the “resource-explicit reproduction” variant used 1.2, 1.4, and 1.7 times as much memory as the direct-interaction models, respectively.

**Table 1.** Model runtimes, square tiled. Forty measurements of elapsed runtime per tick were collected and averaged for each method at each density and population size and at each of three node-placement densities. For the resource-explicit models depicted in this table, a square tiling of nodes was used. Color bars show the runtime of each model, on a different scale within each column, relative to the slowest runtime within that column.

| Model | Node density | Low density (20 individuals per unit area) |  | High density (200 individuals per unit area) |  |
| --- | --- | --- | --- | --- | --- |
|  |  | 100k individuals<br>Runtime per tick (s) | 1m individuals<br>Runtime per tick (s) | 100k individuals<br>Runtime per tick (s) | 1m individuals<br>Runtime per tick (s) |
| Panmictic | NA | 0.16 | 1.59 | 0.16 | 1.59 |
| Fixed-strength | NA | 3.03 | 38.98 | 17.01 | 233.43 |
| Linear | NA | 3.58 | 45.06 | 23.48 | 302.95 |
| Gaussian | NA | 5.76 | 67.88 | 44.81 | 523.23 |
| Inelastic | 12 | 0.79 | 9.25 | 1.20 | 15.54 |
|  | 25 | 0.96 | 11.02 | 1.32 | 17.40 |
|  | 50 | 1.28 | 14.39 | 1.56 | 20.99 |
| Inelastic "fair" | 12 | 0.97 | 11.30 | 1.35 | 17.70 |
|  | 25 | 1.28 | 15.05 | 1.65 | 21.91 |
|  | 50 | 1.91 | 22.10 | 2.23 | 29.85 |
| Inelastic<br>"resource-explicit<br>reproduction" | 12 | 0.94 | 10.43 | 0.81 | 10.29 |
|  | 25 | 1.34 | 14.86 | 1.15 | 14.71 |
|  | 50 | 2.03 | 22.90 | 1.84 | 22.97 |
| Elastic | 12 | 2.43 | 27.39 | 2.69 | 30.75 |
|  | 25 | 3.52 | 40.70 | 3.65 | 41.50 |
|  | 50 | 5.87 | 65.35 | 5.55 | 65.22 |

A key finding of this analysis (Table 1) is that the resource-explicit models were minimally affected by the density of the population, unlike the direct-interaction models. When simulating the higher-density population, even the slowest of the resource-explicit methods significantly outperformed the fastest direct-interaction spatial model (which, in all cases, was the “fixed-strength” model). When simulating a dense population of 1 million individuals, the “resource-explicit reproduction” variant of the inelastic model (the fastest resource-explicit model in this context) was up to 23 times faster than the fixed-strength model. At lower density, the elastic resource-explicit models ran at around the same speed as the fixed-strength model, or in some cases even more slowly (particularly when node density was higher than carrying-capacity density). The inelastic models were at least twice as fast as the elastic model, however, and were thus markedly faster than the fixed-strength model even at lower density.

The reason that the runtimes for the resource-explicit models do not rapidly increase as density increases is that, given a specified population size, a higher-density population entails a smaller landscape, and thus fewer resource nodes (each of which provides more resources). Thus, the overall amount of processing done remains roughly constant. As population density decreases, however, even the most efficient of the resource-explicit models would eventually perform more poorly than a model regulated by direct interactions. The exact density at which this occurs depends on the node density and whether competition is modeled as regulating mortality or fecundity (since that choice affects the size and density of the population when competitive interactions are evaluated).

Very little runtime difference was observed between the square and hexagonal tilings (Supplemental Table 1), while the models in which nodes were randomly distributed during each tick of the model ran at about the same speed at higher population density, but somewhat slower at lower population density (Supplemental Table 2).

Comparing the inelastic model and its two variants, the “fair” variant added only a modest amount to the runtime of the default inelastic model. The “resource-explicit reproduction” variant was slower than the default inelastic model at lower population density, but ran faster at higher density. This was therefore the fastest model in the simulations with higher density, whereas the default inelastic model was the fastest at lower density.

Finally, the panmictic model provides a baseline comparison that shows that the vast majority of the runtime of the other models was indeed spent handling spatial interactions. In models with complex genetics or other computational overhead, spatial calculations may account for a lower proportion of the total runtime. For such models, switching to a resource-explicit modeling approach would thus not yield as great a degree of relative speedup.

## DISCUSSION

In this manuscript, we introduced a resource-explicit modeling approach in which the resources available to each individual are directly represented. Individuals therefore compete only indirectly with one another by interacting with the resources around them. This method comes with several advantages over commonly used direct-interaction spatial models, with runtime improvement arguably being the most salient. The resource-explicit approach will allow for the simulation of hitherto impractically large populations, while also allowing for studies of smaller populations to be conducted more quickly and efficiently. In addition to runtime advantages, there are a number of model features that can be easily and efficiently implemented in a resource-explicit model, some of which are difficult to implement in direct-interaction models. The following six examples provide an incomplete list of such features.

### 1. Heterogeneous landscapes and irregular area boundaries

The resource-explicit approach elegantly lends itself to the simulation of competition on landscapes with heterogeneous resource availability. To accomplish this, nodes can simply be parameterized with different resource amounts depending on the local habitat quality in different areas of the landscape. When an individual forages, the quality of the habitat in which it lives will thus be reflected in the amount of resources available in the nodes that make up its foraging area without any additional computational overhead or additional code (aside from the initial parameterization of the nodes). This holds even for individuals that abut irregular boundaries such as coastlines.

Randomly distributing the resource nodes has notable applicability in modeling heterogeneous landscapes. A model in which nodes are randomly placed at the beginning of the model and then not re-shuffled could be used to generate a new random heterogeneous landscape in each simulation. Alternatively, the version presented in this manuscript, in which nodes are shuffled each tick, could be used in conjunction with a landscape map to allow simulations to run without any pre-calculation of resource-node positions or resource values. In such a model, the landscape map could control the resource values of randomly placed nodes based on their locations, or could be used to implement a more complex node placement algorithm in which nodes are more likely to be placed in higher-quality habitat, using a technique such as rejection sampling. However, grid placements of nodes with precalculated resource values may be preferred for performance reasons, or to minimize variance in interaction strengths.

### 1. 2. Multiple types of resources

Many species require different types of resources at different stages of life, or need a different resource to reproduce than they normally forage on. Some examples include monarch butterflies, which can feed on nectar from many plants but only lay their eggs on milkweed, or mosquitos, which require water habitat resources as larvae, feed on nectar as adults, and only lay eggs after taking a blood meal^9,23,24^. In a resource-explicit model, nodes can be parameterized with a separate value for each type of resources, allowing for the simulation of complex life histories while adding little overhead to the model.

### 1. 3. Interspecies competition

The resource-explicit modeling approach is convenient for implementing models that include competition between multiple species. Each species in such a system need only interact with the resource nodes. By contrast, in direct-interaction models, pairwise interactions as between each species must be considered, resulting in quadratic growth in the complexity of the model. Predator–prey interactions can also be defined in a resource-explicit manner, similar to the way in which the mate-search interaction is defined in the “resource-explicit reproduction” model.

Differently sized foraging areas for different species can be implemented by setting different species to forage from different numbers of nodes in the elastic model, or by setting a different foraging radius in the inelastic model. However, this will be most efficient when the size of the foraging areas of the species in the model are within roughly an order of magnitude of one another, to keep the number of nodes interacted with by each individual within reasonable bounds.

### 1. 4. Resource variability as a function of time

Resource-explicit models are also amenable to simulating variation in resource availability as a function of time, whether in the form of periodic variation (such as seasonality), random variation (such as tree masting), or long-term trajectories (such as changes caused by climate change)^25–27^. To accomplish this, resource nodes can be replenished according to an appropriate function of time, rather than always replenishing to a fixed value.

### 1. 5. Species-induced habitat change

In addition to competing for resources, many species also have a direct effect on the landscape in which they live. For example, many species of bacteria and yeast, such as the beloved *Saccharomyces cerevisiae* (brewer’s yeast), ingest sugars and excretes ethanol^28^. As the concentration of alcohol increases and the availability of sugars decreases, the habitability and the resource availability of their environment both decrease. A resource-explicit modeling approach could allow for the spatial modeling of these processes by explicitly tracking the amount of sugar and alcohol at each resource node, enabling the elegant modeling of spatial environments such as Petri dishes. Some animals can also induce environmental changes that are helpful to themselves (“niche construction”)^29^. A resource-explicit model could, for example, track the increasing abundance of fish in a pond after a beaver extends its dam by changing the resource values at corresponding nodes, and could even simulate the flooding of the landscape using elevation values associated with each resource node.

### 1. 6. Imperfect resource regeneration rate

In many natural systems, groups of animals extract local resources much faster than those resources can replenish. For example, herds of grazing animals might eat grass in their local area much faster than it grows, but the animals survive by moving across the landscape, leaving previously exploited areas time to recover. Such dynamics can be implemented in a resource-explicit model by parameterizing nodes with an appropriate maximum value along with a regeneration function. Such a model could be further enhanced by designing dispersal within the model to be influenced by resource availability such that individuals tend to disperse towards areas where resources are the most plentiful. Another enhancement could be made by incorporating a functional response by consumers to resource density by imposing a penalty on resource collection from low quality areas or areas where resources have not been fully replenished, reflecting the increased effort of finding food when it is sparse. This design would create a strong foundation for a highly detailed model of herd-animal behavior.

To illustrate the possibilities for extending and modifying the basic resource-explicit approach, as discussed above, we provide three additional extensions in the supplement. The first is a variant of the elastic model that is optimized for infrequent dispersal (Additional Extensions part I in the supplement). The second is a variant with a semi-fixed population size, which may be desirable for modelers seeking to construct spatial analogs of analytical models with fixed population sizes (Additional Extensions part II). The third is a scaled-up demonstration model that simulates a population of 10 million rodents on the South Island of New Zealand and which including seasonality and a heterogeneous landscape, yet still achieves good performance.

In the big picture, the resource-explicit modeling approach is related to approaches used in several other fields that utilize spatial approximations or discretizations to increase performance. In computational fluid dynamics, fluids are treated as comprising many discrete cells, since the simulation of individual molecules is not tractable for most applications^30^. In computational astrophysics, the Barnes–Hut method reduces the complexity of *n*-body simulations from *O(n*^2^) to *O(n log n*) by using an octree data structure and then aggregating the interaction forces exerted by distant objects into a combined force vector^31^. As in these other fields, the resource-explicit approach entails some degree of approximation in order to achieve performance gains.

In the case of the resource-explicit approach, this approximation involves exchanging a strict functional correspondence between distance and interaction strength for a system in which interaction strength has a distribution of possible values for a given distance between individuals (Figure 3). The circle-intersection interaction function which this distribution approximates can be biologically rationalized as reflecting resource-foraging behavior, and the indirect interaction strengths in the resource-explicit models match the circle-intersection function quite closely. In many cases, the variation in pairwise interaction strengths is negligible – averaging out across the numerous interactions an individual experiences (Figure 4), or otherwise outweighed by the broader stochasticity of mortality in most individual-based models. In cases where minimizing the variation in interaction strengths is worth slower runtimes, a higher density of resource nodes can be used, whereas a lower density can be used when increased variation is a worthwhile tradeoff for faster runtimes.

There are some scenarios in which the resource-explicit approach might not be the preferred choice. This approach emulates indirect competition between individuals mediated by shared resources. For systems where direct or “interference” competition is more important, it may make more sense to simulate direct interactions between individuals, even at the price of longer runtimes.

Furthermore, in systems with very low densities or small populations, a resource-explicit model might run more slowly than a direct-interaction model. However, runtimes are not the only consideration in model design, and the other properties and capabilities of resource-explicit models may prove desirable even at the cost of runtime speed when modeling sparse populations.

In many contexts, the resource-explicit methods presented here provide excellent performance and a high degree of flexibility. These methods enable spatial simulations of populations that are intractably large to model with direct individual–individual interactions, and facilitate a highly efficient approach to implementing landscape heterogeneity and complex resource-exploitation behaviors. We therefore believe that these new methods will prove an essential addition to the spatial modeler’s toolkit.

## DATA AND CODE AVAILABILITY

The SLiM code for the models and the data used in this manuscript are available at https://github.com/MesserLab/ResourceExplicitModels. The SLiM simulation software used in the project is available at https://messerlab.org/slim/ or at https://github.com/MesserLab/SLiM. Modelers seeking support in developing resource-explicit models based on these models are invited to ask for support by raising issues on the resource-explicit models GitHub page.

## ACKNOWLEDGEMENTS

This study was supported by funding from the National Institutes of Health awards R01GM127418, R01HG012473, and R35GM152242 to P.W.M.

## Supplemental Information

## RESOURCE NODE PLACEMENT METHODS

Three resource node placement methods were used in the models assessed in this manuscript. These methods were: placing resource nodes at the center of each square in a uniform square tiling of the landscape (Figure 2), placing resource nodes at the center of each hexagon in a uniform hexagonal tiling of the landscape (Supplemental Figure 1), and placing the resource nodes randomly across the landscape, re-randomizing the placement of the nodes during each tick of the model (Supplemental Figure 2). Specifically, random placement was accomplished by assigning each node an x and y coordinate randomly drawn from a uniform distribution with a minimum of 0 and a maximum of the side length of the model. For each of these placement methods, several resource node densities were tested.

In terms of code complexity, the randomized placement method is the simplest. This method allows resource nodes to be added to the model with just a few lines of code. The code to perform square and hexagonal tilings in our models is significantly more complex due to the desire for the tiling to cover any square area of arbitrary dimensions. To accomplish this, the grid of nodes is centered within the modeled area, and nodes with areas that fall partially outside the landscape are parameterized with a proportionately decreased amount of resources. For example, if tiling a 2.8 by 2.8 landscape with square nodes that are each 1 by 1, a 3 by 3 grid of nodes would be used, with the grid of nodes centered within the area, and with nodes along the edges of the area are parameterized with an appropriately reduced amount of resources given the portion of the node’s area that falls within the modeled area. In the hexagonally tiled models, the code required to accomplish this task is somewhat more complex than in the square-tiled models. That said, this complexity was included in the interest of matching the direct-interaction models in this manuscript as closely as possible. In models designed from the ground up as resource-explicit that do not need to match a prior model, the landscape could be defined as having a boundary exactly matching a desired grid of nodes, removing the need for this complexity. Minor adjustment to these methods would also be appropriate if modeling a periodic area (such as a toroidal landscape), in order to ensure appropriate resource availability along the “seams” of the model.

### Choice of node density and placement method

As demonstrated in the results section of this manuscript, the choice of resource node density effects the runtime of the model as well as the magnitude of variance in interaction strengths within the model. An increased node density tends to yield a tighter distribution of interaction strengths, but runs more slowly. The choice of node density must therefore be made based on what kind of interaction strength variance can be can be considered acceptable and how fast the model needs to run.

The choice between a regularly tiled model or a model with randomly distributed nodes is also fairly straightforward. Models with regular tilings have tighter distributions of interaction strengths, and run slightly faster compared to models where the node positions are re-randomized during each tick of the model. However, a model with randomized resource positions that are not re-randomized could be used to represent a random heterogeneous landscape that is uniquely generated each time the model is run. Random node placement could also be combined with a landscape map to allow for the simulation of a specified heterogeneous landscape without any pre-calculation step. Additionally, more complex node placement strategies may provide desirable ways to represent realistic resource variability. For example, nodes might be placed according to a Poisson-Disc Sampling, in which entities are randomly placed, but are not placed closer to one another than a specified minimum distance.

When minimizing variance in interaction strength is a priority, a regular tiling of resource nodes is the best option (Supplemental Figure 3; compare Figures 3 and 4 versus Figure 5). This holds true both for simulations that are intended to emulate a homogenous space (as is common in direct-interaction models) and for simulations involving heterogeneous landscape maps (in which case pre-calculating the appropriate amount of resources at each node and using a dense tiling will result in the resource-explicit model matching the landscape as closely as possible).

The choice between a hexagonal grid and a square grid is less consequential. The two are almost interchangeable. Since a square grid is simpler to implement, it may be the right choice for most models. In most cases, the standard deviation in interaction strengths as compared to the circle-intersection function with one tiling or another at a given density varies by no more than a percent or two (Supplemental Figure 3).

There are only three possible ways to tile a plane with edge-to-edge congruent regular polygons: using equilateral triangles, squares, or hexagons^12,32^. In other ecological modeling contexts, such as modeling an area by using an array of panmictic demes linked by migration, it is well accepted that spatial artifacts are smaller when using hexagonal grids compared to using square grids^12^. Without going into complete detail, the reason the tiling does not matter as much in the resource-explicit method is as follows: in our method, the *area* of each shape (be it a hexagon or square) is determined by node density parameter. In linked panmictic models, the *distance* between nodes is fixed by the model (with the distance between what are considered to be different populations being related to the dispersal characteristics of the species being modeled). Given a fixed distance between nodes, a hexagonal tiling is a denser packing, with each hexagon therefore having a smaller area than squares would have in a square tiling with the same distance between nodes. Thus, a hexagonal tiling offers a large advantage in linked panmictic models, but is not that different from a square tiling in our resource-explicit models. However, in some cases, a hexagonal tiling has some desirable properties in terms of minimizing unequal availability of resource nodes in the inelastic model (see Supplemental Figures 14-19).

## DYNAMICS OF THE RESOURCE-EXPLICIT MODELS

In the results section, interaction strengths in the resource-explicit models were assessed by measuring pairwise interactions between randomly placed individuals. The interaction strengths measured in the resource-explicit models were then subtracted from the strengths measured by the circle-intersection function. This analysis was performed using a square tiling of the landscape (Figure 4), a hexagonal tiling of the landscape (Supplemental Figure 4), and with resource nodes randomly placed, with new positions drawn for each interaction measured (Supplemental Figure 5).

A further analysis was performed that compared the overall interaction forces experienced by individuals in the resource-explicit models to that calculated by the circle-intersection function. This analysis was performed using a square tiling of the landscape (Figure 5), a hexagonal tiling of the landscape (Supplemental Figure 6), and with resource nodes randomly placed, with new positions drawn for each interaction measured (Supplemental Figure 7).

We performed an addition assessment of interaction strengths by measuring an “interaction field” between a focal individual and a dense grid of points within interaction range. This analysis was performed with a focal individual placed at a coordinate that minimized the variance between the resource-explicit model and the circle-intersection function, and again at a coordinate that maximized the variance. This analysis was conducted with a hexagonally tiled landscape as well as a square-tiled landscape, each at a density of 12, 25, and 50 nodes per unit area, each with both the inelastic and elastic method (Supplemental Figures 8-13).

### Unequal Resource Availability in the Inelastic Model

In the inelastic implementation of the method, individuals are not guaranteed to forage from the exact nominal foraging area – an individual might forage from more or fewer nodes. When the area is tiled with a uniform grid of nodes, the exact position of an individual relative to this tiling is what determines how many nodes fall within its foraging radius (Supplemental Figures 14-19).

## COMPUTATIONAL PERFORMANCE OF THE RESOURCE-EXPLICIT MODELS

The runtime differences between the square-tiled models and hexagon-tiled models were negligible, representing at most a small fraction of the runtime of the model (Supplemental Table 1). When resource nodes were randomly distributed during each tick of the model, runtimes were slower (Supplemental Table 2). Unsurprisingly, the models most slowed were those with the most resource nodes, which in some cases took almost twice as long to run as a uniformly tiled model with the same population size and node density.

## ADDITIONAL EXTENSIONS

### I. An Implementation of the Elastic Method Optimized for Infrequent Dispersal

An improvement in runtime can be achieved in the elastic model if each individual only disperses infrequently or only a single time (their initial dispersal from their maternal parent), by finding the nearest resource nodes to an individual only after it disperses, and then caching and reusing this list of nodes during each subsequent tick until the individual disperses again or dies. Individuals disperse only once in all of the models presented in this manuscript, but this optimization was not made in the default elastic model in order to provide an accurate reflection of the runtime that could be expected in a model where individuals move during each tick. This optimization is not relevant in a model with non-overlapping generations or in a model with nodes that are re-randomized each time step, since the cached node lists would never be reused.

The degree of runtime improvement that this optimization could yield is highly dependent on the demography of the model. In models with a low per-tick mortality and in which individuals only disperse once, this method could be faster than all of the other resource-explicit methods presented in this manuscript. In the model assessed in this study, the mortality rate was quite high (about 80 percent per tick) due to the large number of offspring produced every tick of the model, and the cached node lists only save time for individuals that survive to the next tick to use that cache. Thus, when modified to include this optimization, the elastic model used in this study only increased in speed by a small amount, and was still much slower than the inelastic model. The full code for this modification is provided on GitHub.

### II. A Resource-Explicit Model with a Semi-Fixed Population Size

Many analytical models consider populations to consist of a fixed number of individuals. When extending pre-existing analytical models into an individual-based spatial context, it can be desirable to otherwise match the analytical model as closely as possible by maintaining a fixed population size in the spatial model (recognizing that this is biologically unrealistic). A modification of the resource-explicit modeling technique allows for a population size that is fixed except in the event of extreme disruptions to the population.

In implementations described in the methods section, each node is parameterized with some amount of resources that depends on the density of the modeled species and the density of the resource nodes, and each node distributes its resources evenly to all individuals that forage from it. If, for example, there are 50 individuals foraging from a node with 10 resources available, each individual will receive 0.2 resources, contributing a 20 percent probability of survival to each of those individuals.

In this variant of the model with a semi-fixed population size, resource nodes are instead parameterized by an integer number of “tickets”. Instead of receiving a floating-point amount of resources, individuals instead have a chance to receive a ticket. After tickets are distributed, only individuals who have received a ticket survive. This results in the landscape maintaining a population of exactly the specified carrying capacity, except in the event of a disturbance in the model that causes reproduction to be insufficient to reach that carrying capacity (such as a simulated intervention against a pest population). Note that the population will also not stay exactly at a defined capacity in a model with a marginal habitat quality or a very low birthrate, in which case reproduction may sometimes produce fewer individuals than mortality removes even when no external factors are present.

For each of the four variants of the resource-explicit model presented in this manuscript (elastic, inelastic, inelastic “fair”, and inelastic with “resource-explicit reproduction”), the GitHub file repository for this project also contains an equivalent model that has been modified as described above to maintain a semi-fixed population size. This modification results in a moderate performance reduction in the elastic models, but does not appear to noticeably affect the performance of the inelastic models.

For modelers seeking to design spatial models that match analytical models as closely as possible, we anticipate that the “fair” variant of the inelastic model may be the best option. This model is faster than the elastic model, yet it avoids the small-scale spatial artifacts present in the default inelastic model which may be undesirable in this context (see Figure 4 and Supplemental Figures 14-19).

### III. A Spatial Model of the South Island of New Zealand

The resource-explicit method has the potential to scale up to modeling large populations in large heterogeneous habitats while maintaining relatively performant runtimes. As an initial exploration into this possibility, we implemented a model on a landscape map of the South Island of New Zealand. Endemic diversity in New Zealand is threatened by the presence of numerous invasive mammal species, and detailed spatial modeling is the first step in investigating potential population control strategies, such as gene drive, in order to maintain and restore biodiversity^33^.

To produce a heterogeneous landscape map of the island, we chose to define habitat quality as a function of elevation, with optimum habitat at about 300 meters above sea level, with quality decreasing at higher and lower elevations. Each resource node was parameterized with resources according to the local elevation near that node. The total area of the South Island is 150,416 km^2^, but mountainous regions were considered to have no accessible resources^34^. The amount of the landscape defined as usable habitat by the generic focal species in the model was 100,636 km^2^. The nominal foraging area of the focal species was defined as 0.25 km^2^, and the landscape was populated with nodes using a square tiling at a density of 12 nodes per 0.25 km^2^. The total number of resource nodes is 4.8 million. Ticks in the model represent monthly intervals. We included seasonality in the model as a per-tick multiplier to the resource value of each node that follows a sinusoidal function with a maximum of 1 in the summer and 0.5 in the winter. The carrying capacity of the focal species across the entire area is 10 million individuals in the summer, with a maximum density of about 50 individuals per 0.25 km^2^ and an average density of about 25 individuals per 0.25 km^2^ (and half these densities in the winter). Node coordinates and resource values were determined in a pre-processing step that generated a CSV file that was reused by all of the replicates of the simulation.

The inelastic implementation of the resource-explicit model was chosen for this simulation, in the interest of maximizing the speed of the model. Other than loading node positions from the external CSV file at the outset of the model, the only change to the model was to ensure that individuals were positioned on the landmass at the start of the simulation and were prevented from dispersing into the ocean.

No in-depth analyses were conducted of this model, nor were any analogous models constructed for comparison purposes. The average runtime of each tick of this model was under 1.5 minutes even in the summer, when the population was at its maximum size. This is almost certainly sufficiently fast to be used in a study, unless a very large number of ticks needs to be simulated. Though the dynamics within the model were not thoroughly analyzed, a visual assessment indicates that the heterogeneity of the landscape is satisfactorily reflected in the distribution of the population (Supplemental Figure 20).

**Supplemental Figure 1.**
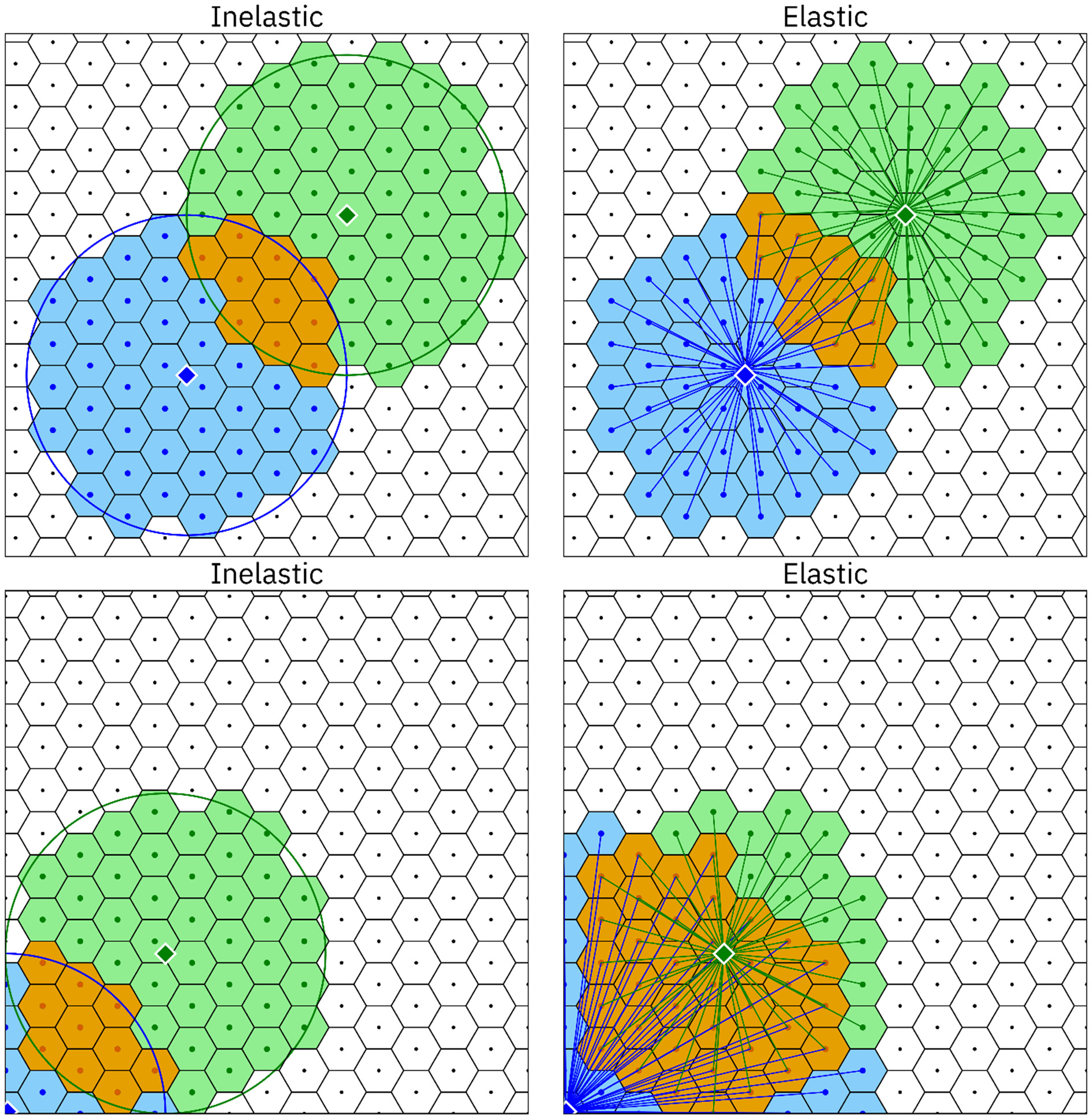
Visualization of resource-explicit interaction algorithms, hexagonal tiling. Competition between two individuals (blue and green diamonds with white outlines) is determined by the portion of their foraging area that overlaps and by the interaction algorithm used in the model. The foraging areas are represented by blue and green shading; the overlapping area is shaded orange. In the inelastic model (left), individuals forage from resource nodes (dots at the center of each hexagon) within their foraging radius. In the elastic model (right), individuals forage from as many nodes as necessary to maintain a nominally sized foraging area (in this case, that area comprises 50 nodes). In upper left panel, the blue individual happens to forage from only 49 nodes, and competition between these two individuals is slightly reduced. In the corner of the landscape, the difference between the two models is much greater: the blue individual has a much smaller foraging area in the inelastic model, while the blue individual in the elastic model forages from much further away to maintain a full-sized foraging area, resulting in greater competition.

**Supplemental Figure 2.**
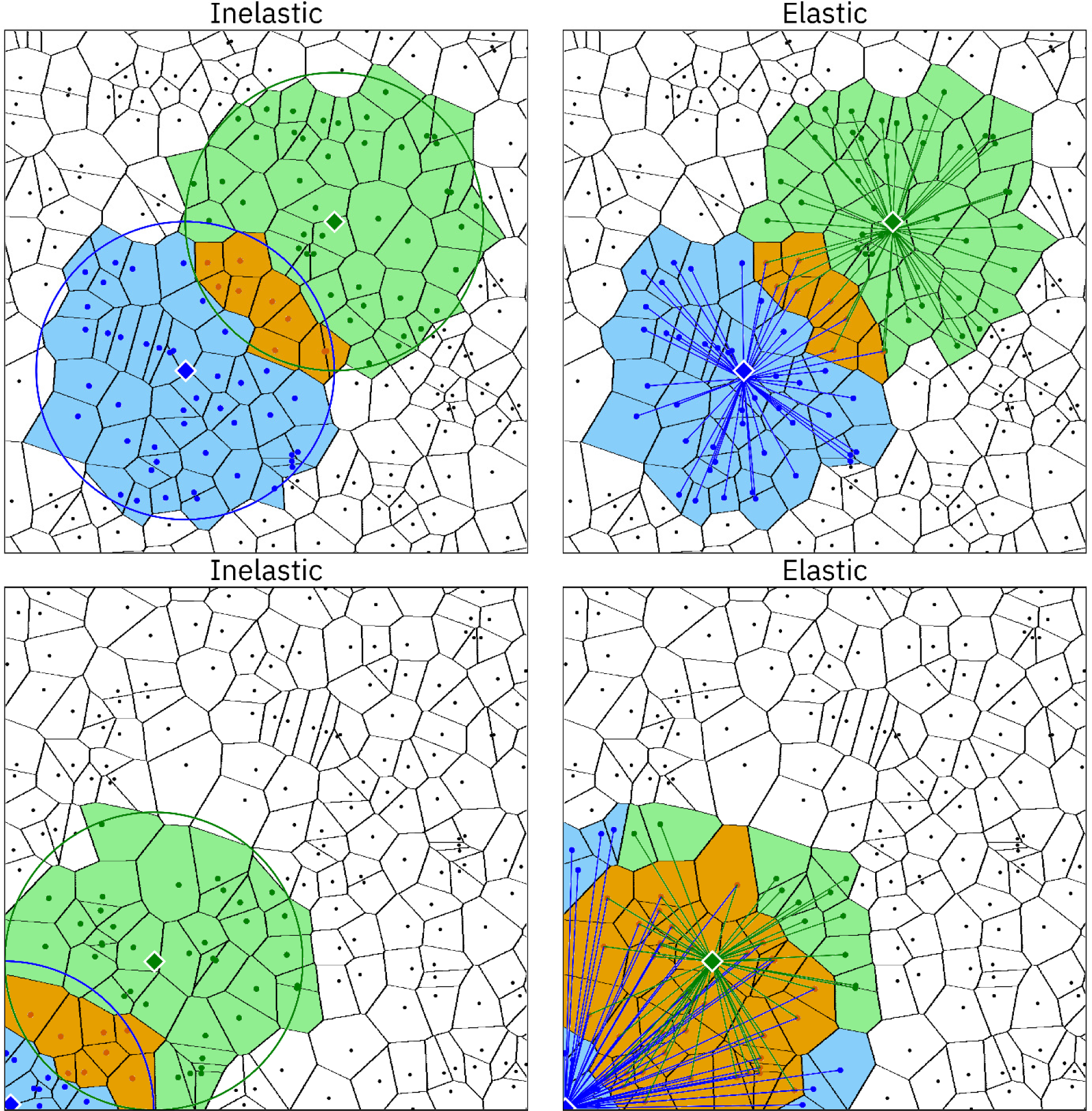
Visualization of resource-explicit interaction algorithms, random node placement. Competition between two individuals (blue and green diamonds with white outlines) is determined by the portion of their foraging area that overlaps and by the interaction algorithm used in the model. The foraging areas are represented by blue and green shading; the overlapping area is shaded orange. In the inelastic model (left), individuals forage from resource nodes (dots at the center of each polygon) within their foraging radius. In the elastic model (right), individuals forage from as many nodes as necessary to maintain a nominally sized foraging area (in this case, that area comprises 50 nodes). When nodes are randomly distributed, the difference between the inelastic and elastic method even in the interior areas is greater. For example, the blue individual in the upper left panel forages from 53 nodes, and the green individual in the bottom left panel forages from 60 nodes, whereas in the elastic model, all individuals forage from 50 nodes, regardless of their position.

**Supplemental Figure 3.**
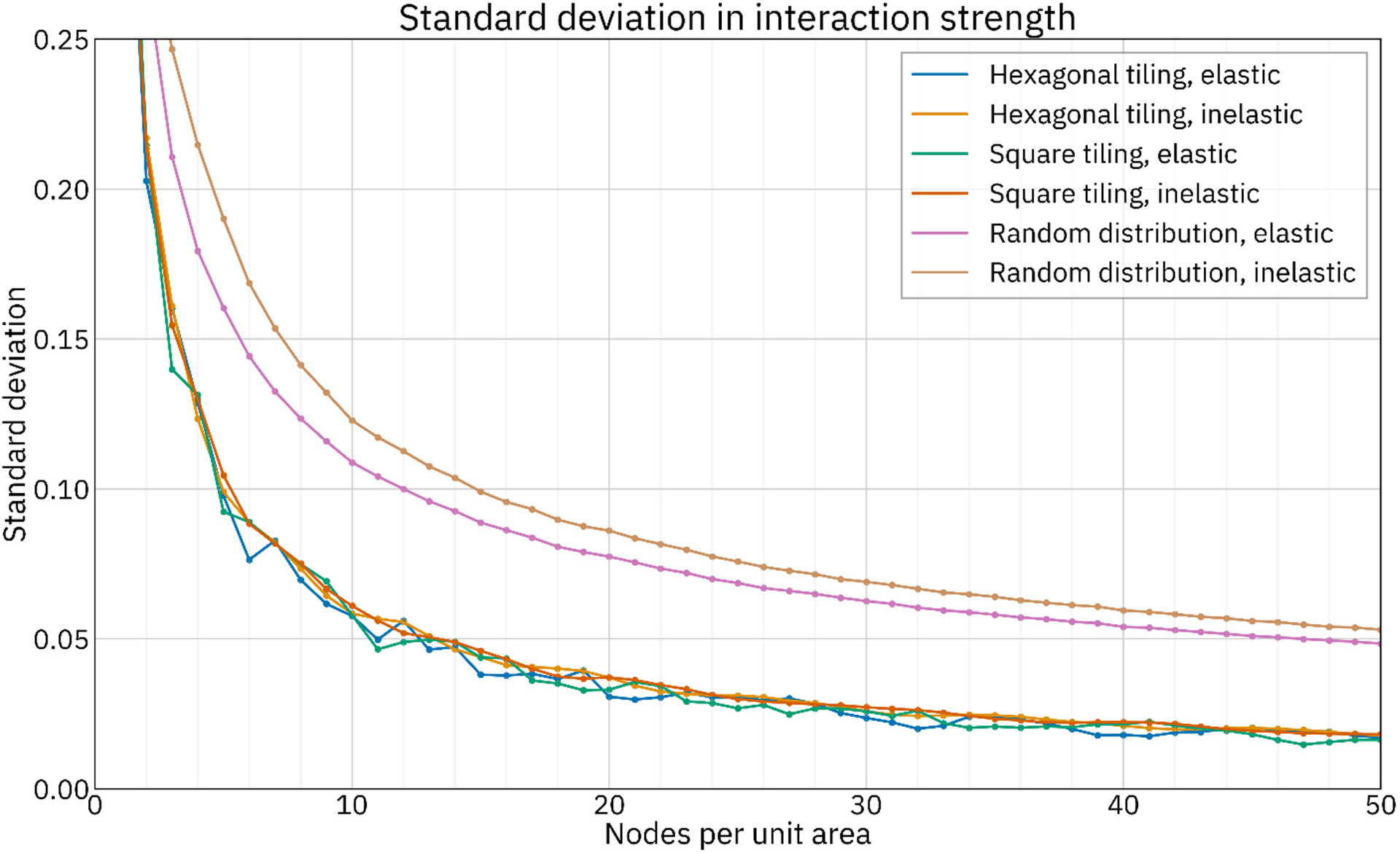
Standard deviation of the differences in interaction strength between resource-explicit models and the circle-intersection function, square tiling. Two million pairwise interaction strengths measured by the circle-intersection function were subtracted from those measured in the resource-explicit models to yield a distribution for each model. The standard deviation was then measured for that distribution. This was repeated with tiling density ranging from 1 to 50 nodes per unit area. Note: while perhaps surprising, the cases where a denser tiling has a higher standard deviation are not due to measurement error (e.g., a hexagonally tiled elastic model with 7 nodes per unit area has a higher standard deviation than the same model with a density of 6 nodes per unit area).

**Supplemental Figure 4.**
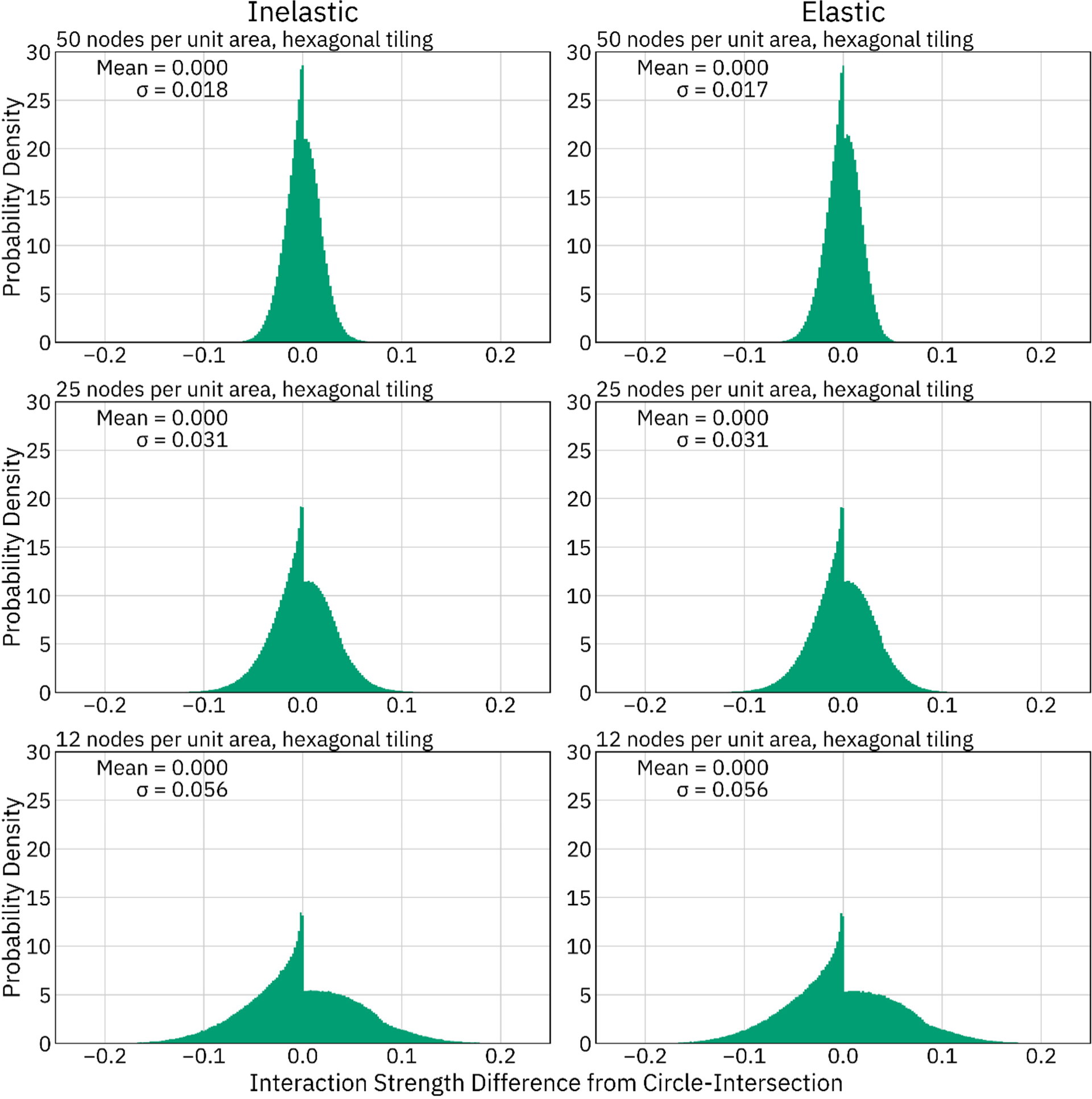
Differences in interaction strength between resource-explicit models and the circle-intersection function, hexagonally-tiled. Two million pairwise interaction strengths between randomly placed individuals as measured by the circle-intersection function were subtracted from those measured in resource-explicit models to yield a distribution of the deviation from the circle-intersection function for each model. As node density increases, the standard deviation decreases. These distributions all have a distinctive peak just below 0 due to cases where pairs of individuals have a very small but non-zero interaction strength when using the circle-intersection function, but the small overlapping portion of their foraging areas does not include any resource nodes.

**Supplemental Figure 5.**
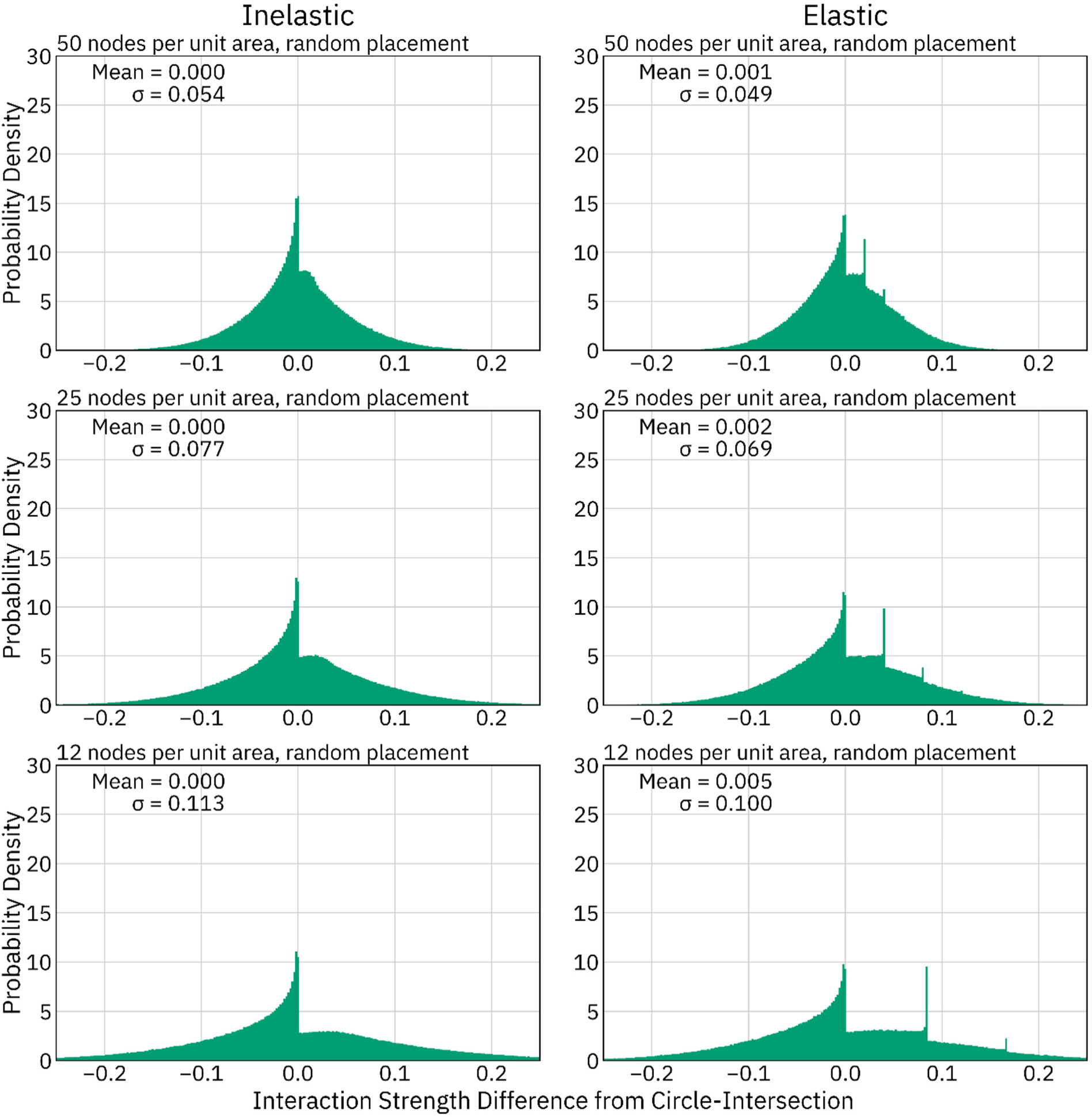
Differences in interaction strength between resource-explicit models and the circle-intersection function, random node placement. Two million pairwise interaction strengths between randomly placed individuals as measured by the circle-intersection function were subtracted from those measured in resource-explicit models to yield a distribution of the deviation from the circle-intersection function for each model. As node density increases, the standard deviation decreases. These distributions all have a distinctive peak just below 0 due to cases where pairs of individuals have a very small but non-zero interaction strength when using the circle-intersection function, but the small overlapping portion of their foraging areas does not include any resource nodes. The distinctive peaks at positive values in the elastic model represent cases where the foraging circles do not intersect at all, but two individuals nonetheless share one or two nodes due to a lack of resource nodes closer to the individuals.

**Supplemental Figure 6.**
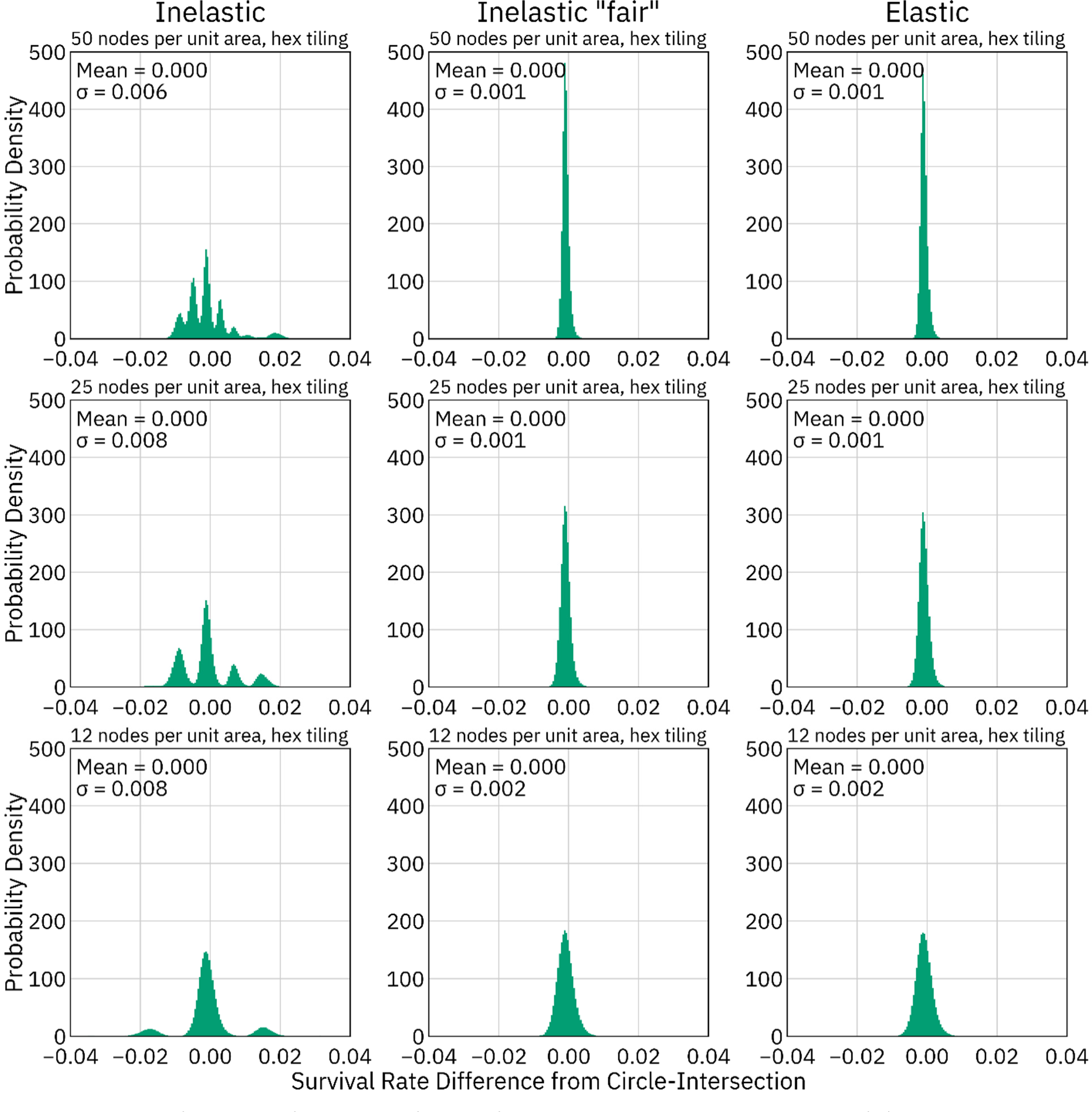
Differences in survival rate between resource-explicit models and a direct-interaction model using the circle-intersection function, hexagonally-tiled. The survival rates of 100,000 individuals were measured using the inelastic method, the “fair” inelastic method, the elastic method, and the circle-intersection function. Strengths measured by the circle-intersection function were subtracted from measurements made using the other resource-explicit methods for each individual to yield distributions of differences.

**Supplemental Figure 7.**
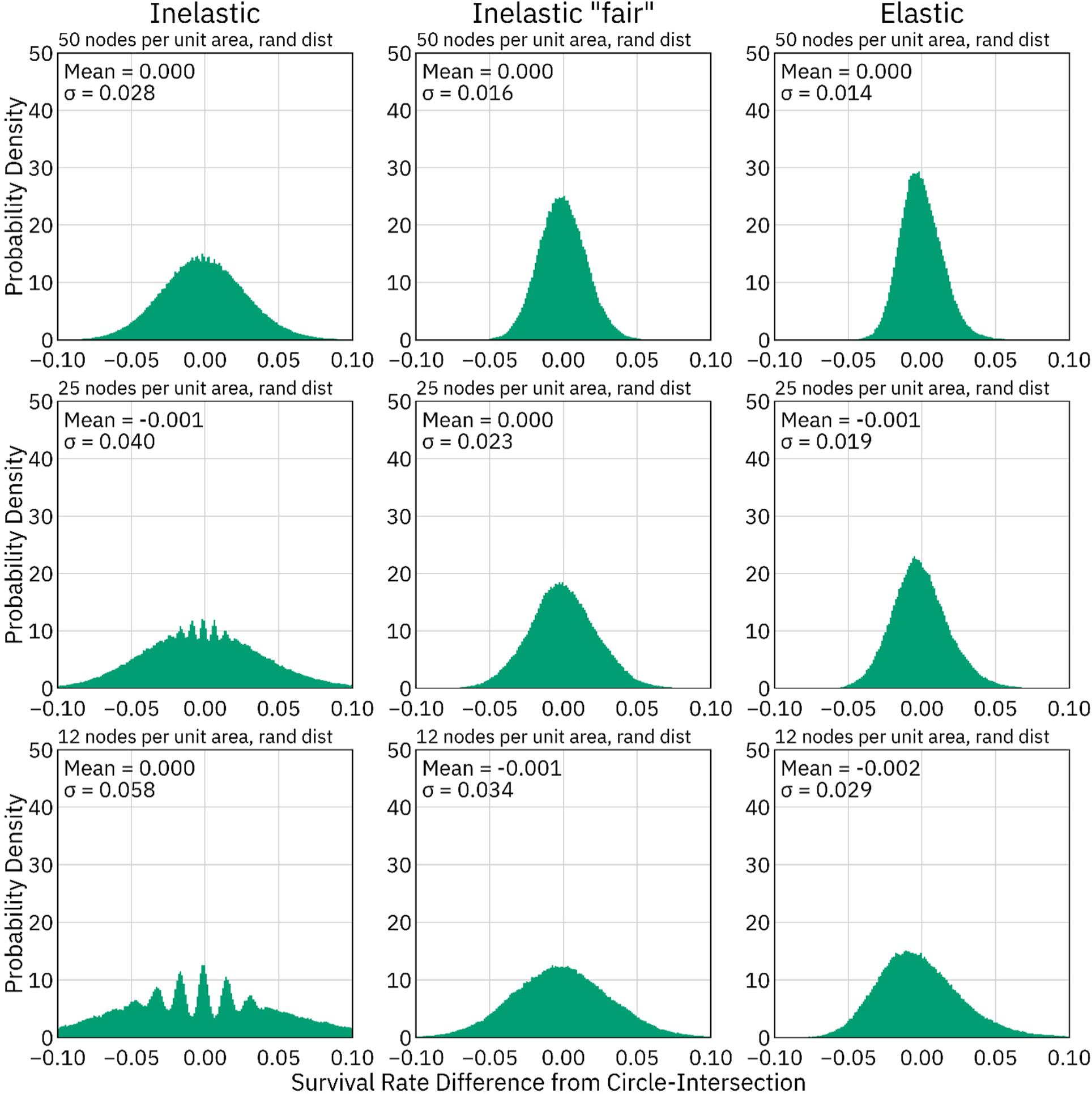
Differences in survival rate between resource-explicit models and a direct-interaction model using the circle-intersection function, random node placement. The survival rates of 100,000 individuals were measured using the inelastic method, the “fair” inelastic method, the elastic method, and the circle-intersection function. Strengths measured by the circle-intersection function were subtracted from measurements made using the other resource-explicit methods for each individual to yield distributions of differences.

**Supplemental Figure 8.**
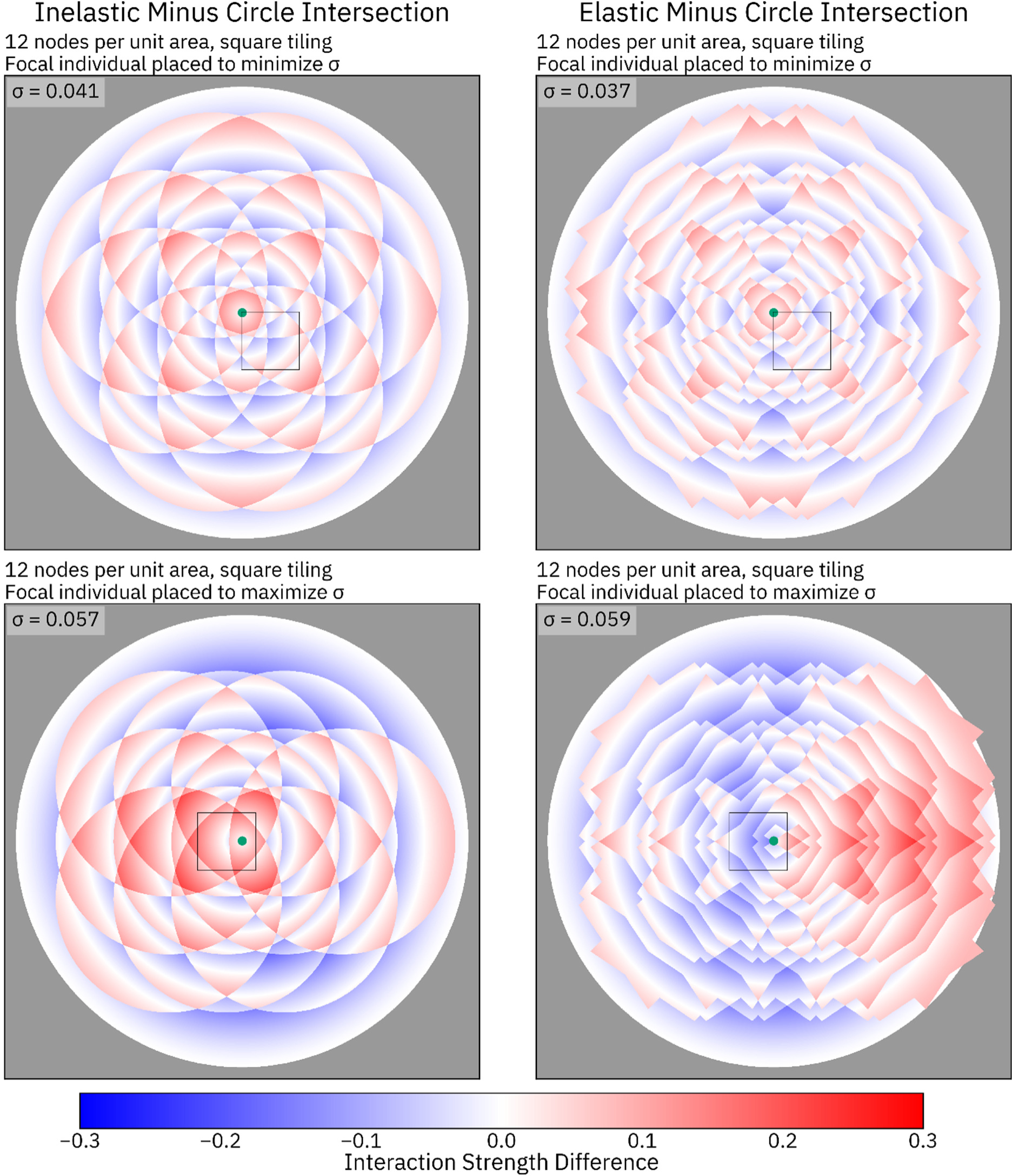
Visualization of the resource-explicit interaction, square tiling, 12 nodes per unit area. A central individual (green dot) was placed to either minimize (top panels) or maximize (bottom panels) the standard deviation between the resource-explicit functions and the circle-intersection function. Interaction strength between the central individual and each other pixel of the image was measured using both a resource-explicit method and the circle-intersection function, and the difference is depicted. Blue shading indicates that the circle-intersection function is stronger, while red shading indicates that the resource-explicit interaction is stronger.

**Supplemental Figure 9.**
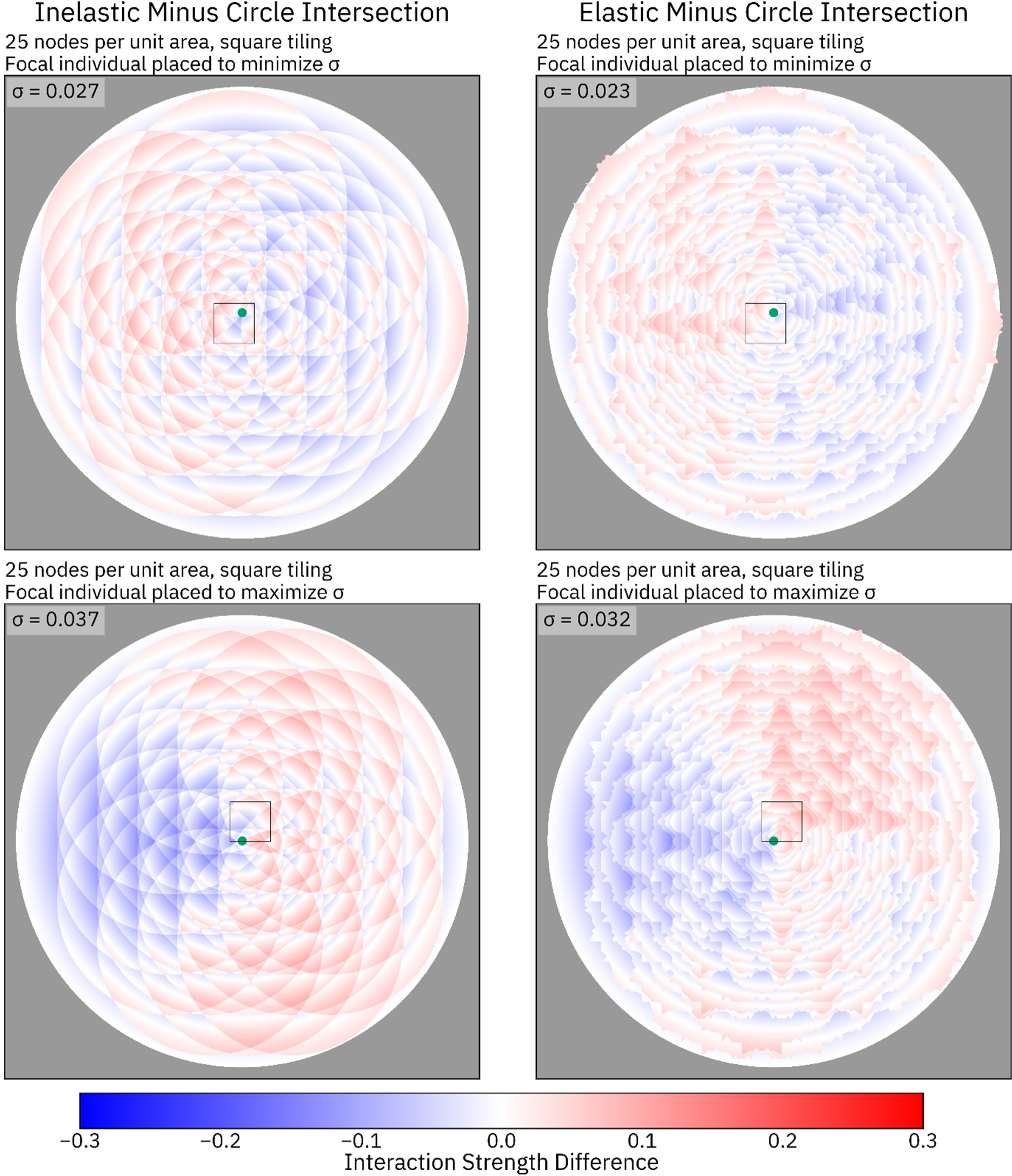
Visualization of the resource-explicit interaction, square tiling, 25 nodes per unit area. A central individual (green dot) was placed to either minimize (top panels) or maximize (bottom panels) the standard deviation between the resource-explicit functions and the circle-intersection function. Interaction strength between the central individual and each other pixel of the image was measured using both a resource-explicit method and the circle-intersection function, and the difference is depicted. Blue shading indicates that the circle-intersection function is stronger, while red shading indicates that the resource-explicit interaction is stronger.

**Supplemental Figure 10.**
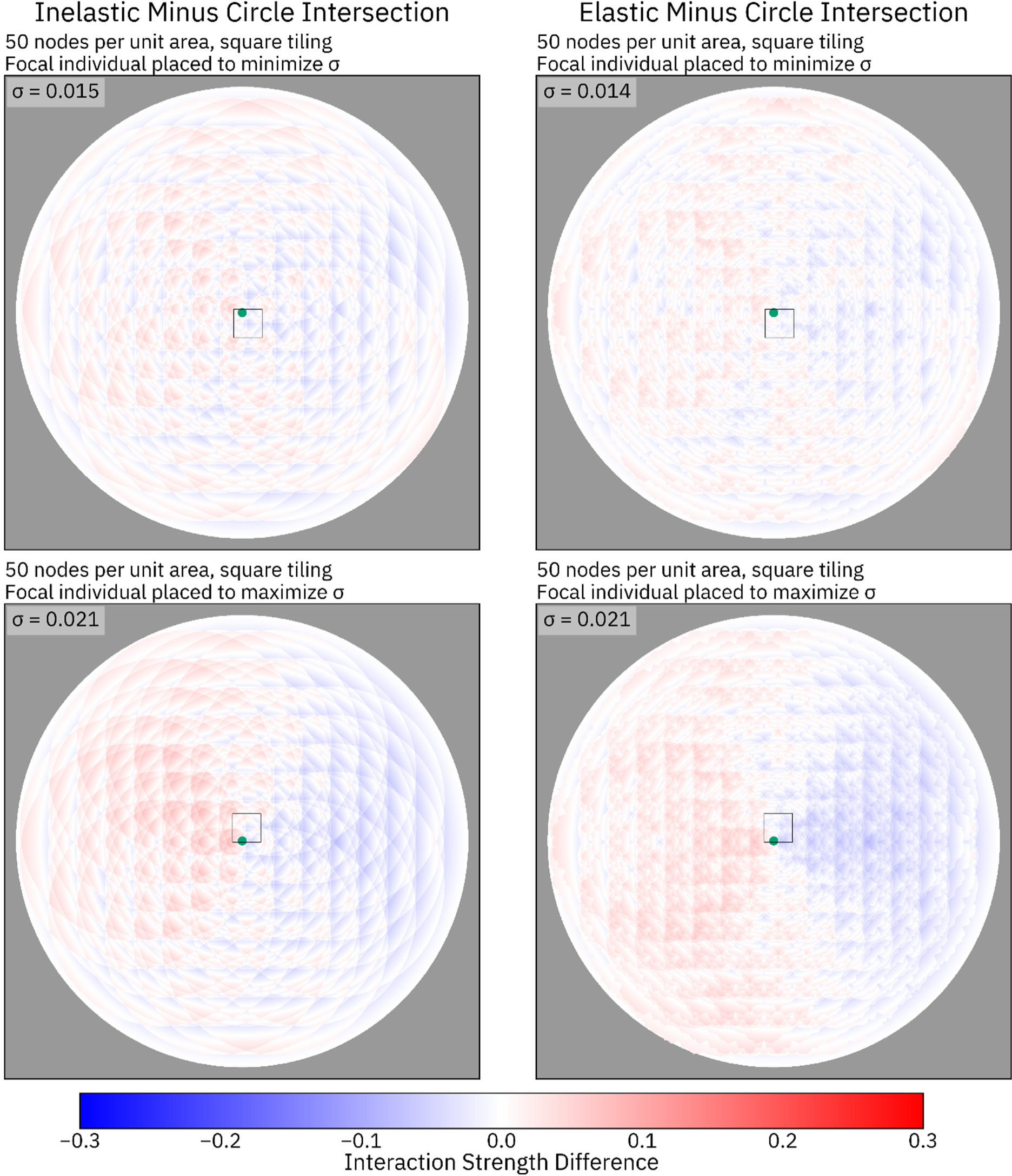
Visualization of the resource-explicit interaction, square tiling, 50 nodes per unit area. A central individual (green dot) was placed to either minimize (top panels) or maximize (bottom panels) the standard deviation between the resource-explicit functions and the circle-intersection function. Interaction strength between the central individual and each other pixel of the image was measured using both a resource-explicit method and the circle-intersection function, and the difference is depicted. Blue shading indicates that the circle-intersection function is stronger, while red shading indicates that the resource-explicit interaction is stronger.

**Supplemental Figure 11.**
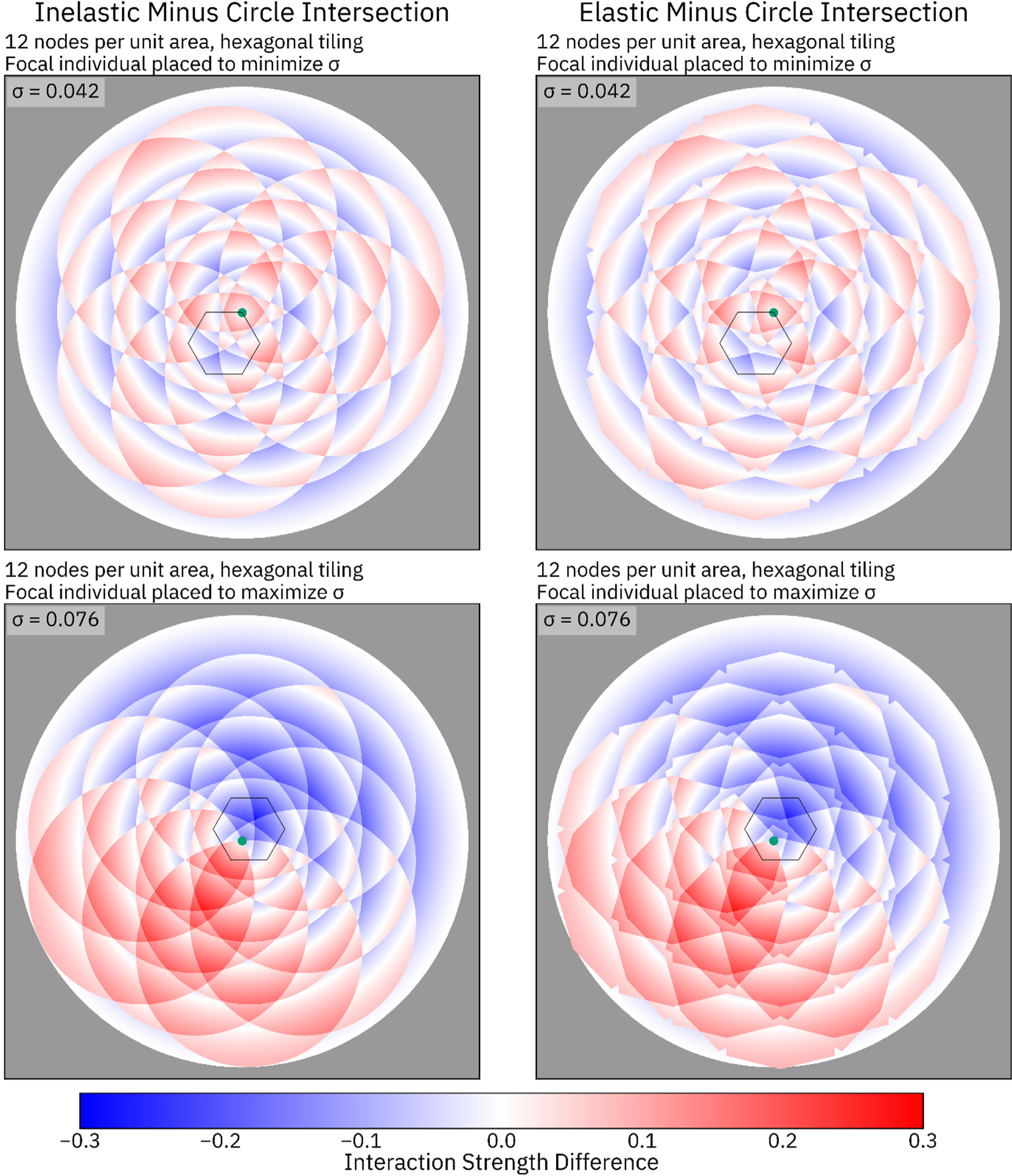
Visualization of the resource-explicit interaction, hexagonal tiling, 12 nodes per unit area. A central individual (green dot) was placed to either minimize (top panels) or maximize (bottom panels) the standard deviation between the resource-explicit functions and the circle-intersection function. Interaction strength between the central individual and each other pixel of the image was measured using both a resource-explicit method and the circle-intersection function, and the difference is depicted. Blue shading indicates that the circle-intersection function is stronger, while red shading indicates that the resource-explicit interaction is stronger.

**Supplemental Figure 12.**
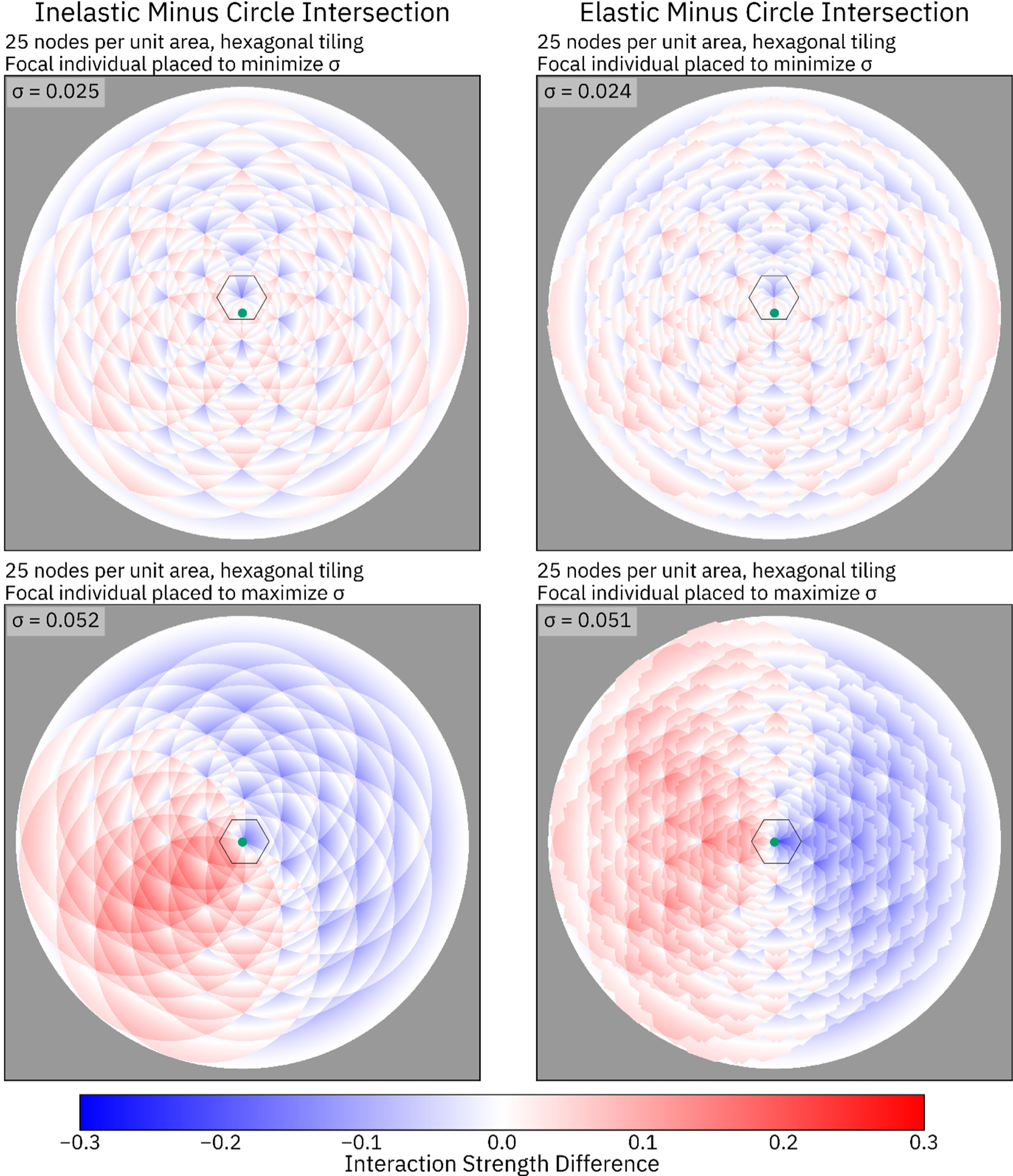
Visualization of the resource-explicit interaction, hexagonal tiling, 25 nodes per unit area. A central individual (green dot) was placed to either minimize (top panels) or maximize (bottom panels) the standard deviation between the resource-explicit functions and the circle-intersection function. Interaction strength between the central individual and each other pixel of the image was measured using both a resource-explicit method and the circle-intersection function, and the difference is depicted. Blue shading indicates that the circle-intersection function is stronger, while red shading indicates that the resource-explicit interaction is stronger.

**Supplemental Figure 13.**
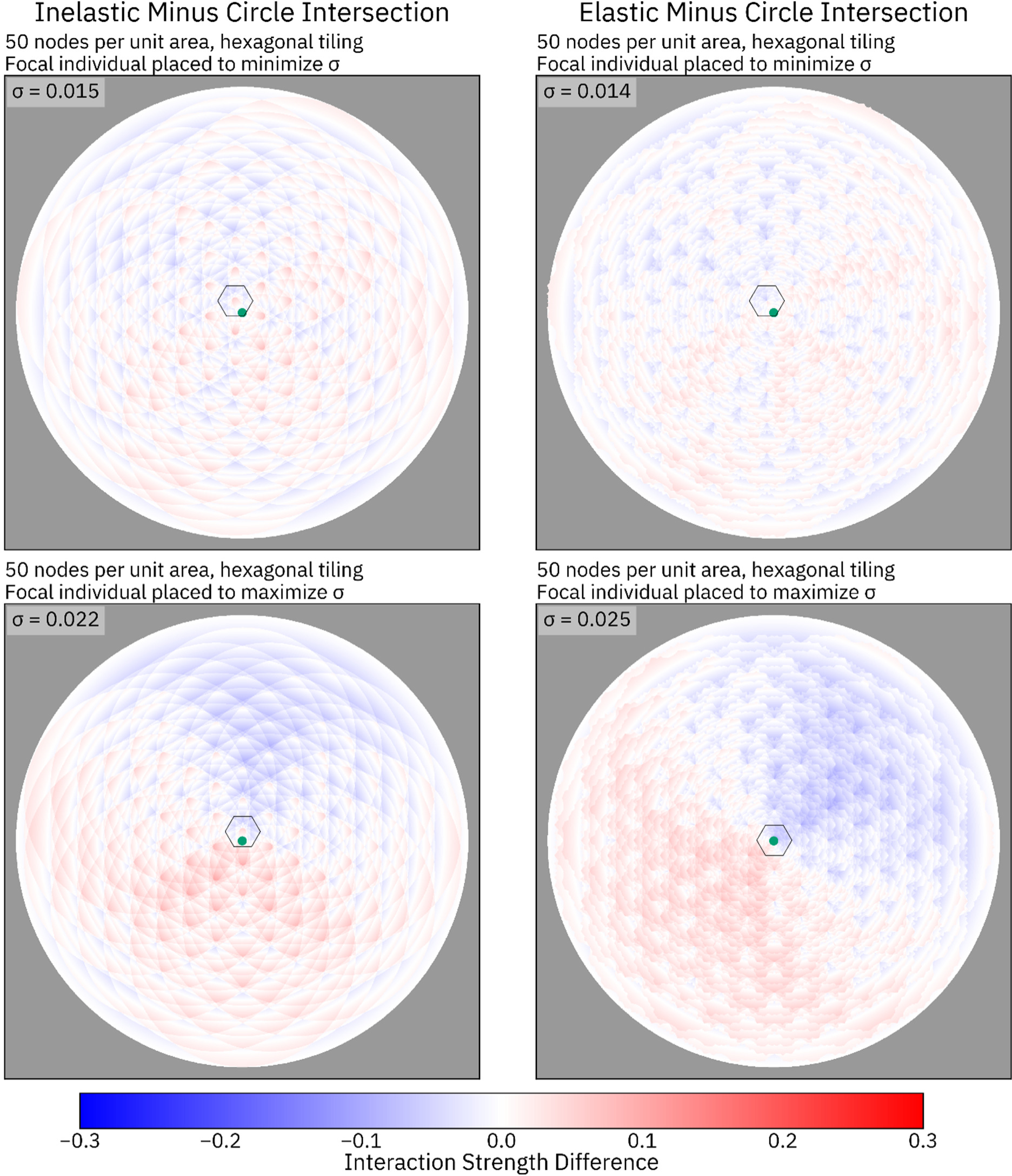
Visualization of the resource-explicit interaction, hexagonal tiling, 50 nodes per unit area. A central individual (green dot) was placed to either minimize (top panels) or maximize (bottom panels) the standard deviation between the resource-explicit functions and the circle-intersection function. Interaction strength between the central individual and each other pixel of the image was measured using both a resource-explicit method and the circle-intersection function, and the difference is depicted by blue and red shading. Unlike in Supplemental Figures 8-12, the coordinate that maximized σ differs between the elastic and inelastic method in this case.

**Supplemental Figure 14.**
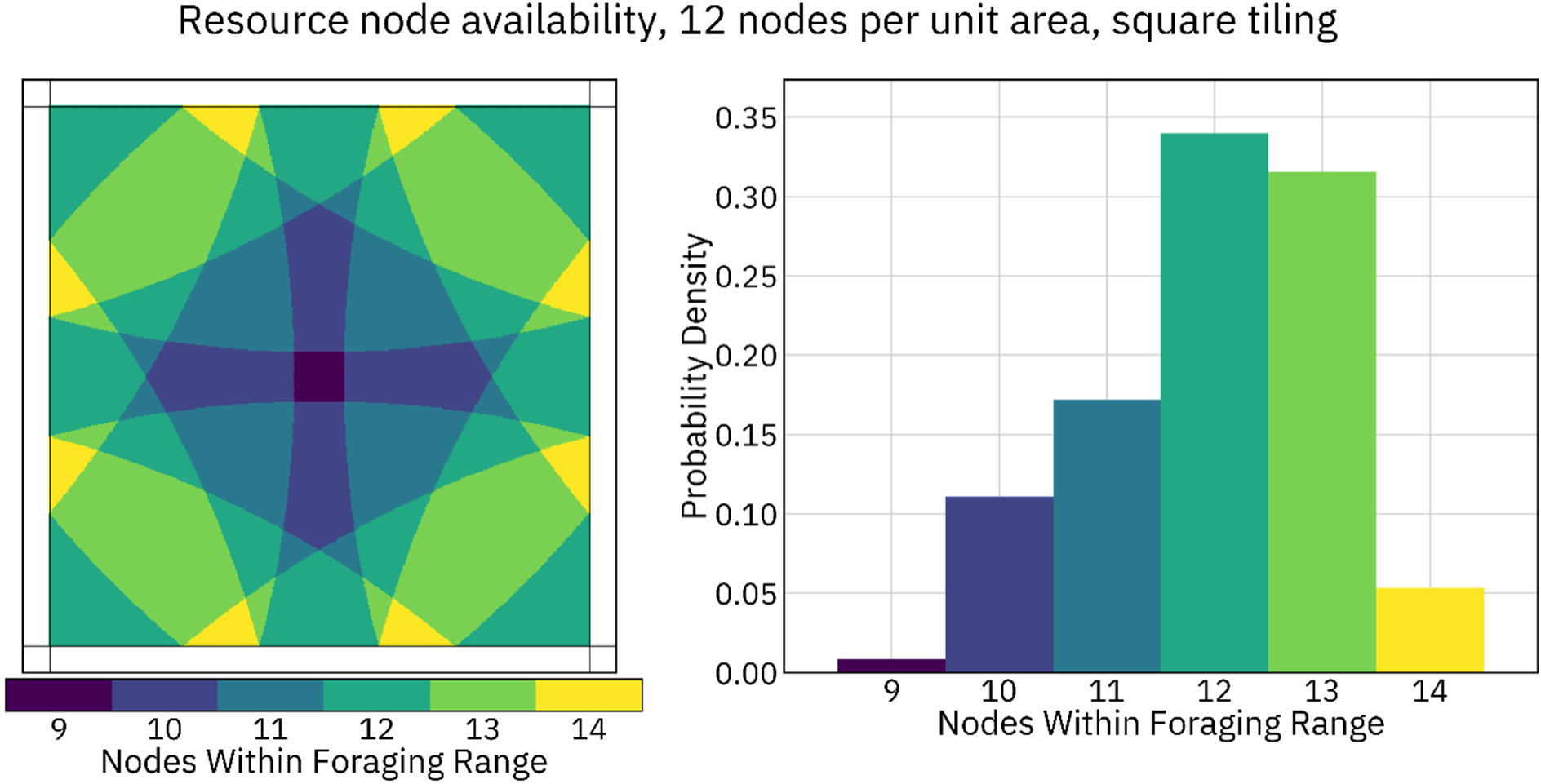
Unequal access to resource nodes, square tiling, 12 nodes per unit area. The number of resource nodes within the foraging radius of an individual depends on their location within a square tile. Individuals located in darker regions of the square (left panel) forage from fewer nodes, while those in brighter regions forage from more. The relative area of each color is depicted in the right panel. The mean value of this distribution is 12.00, and the standard deviation is 1.11 (equivalent to 9.2 percent of the foraging area).

**Supplemental Figure 15.**
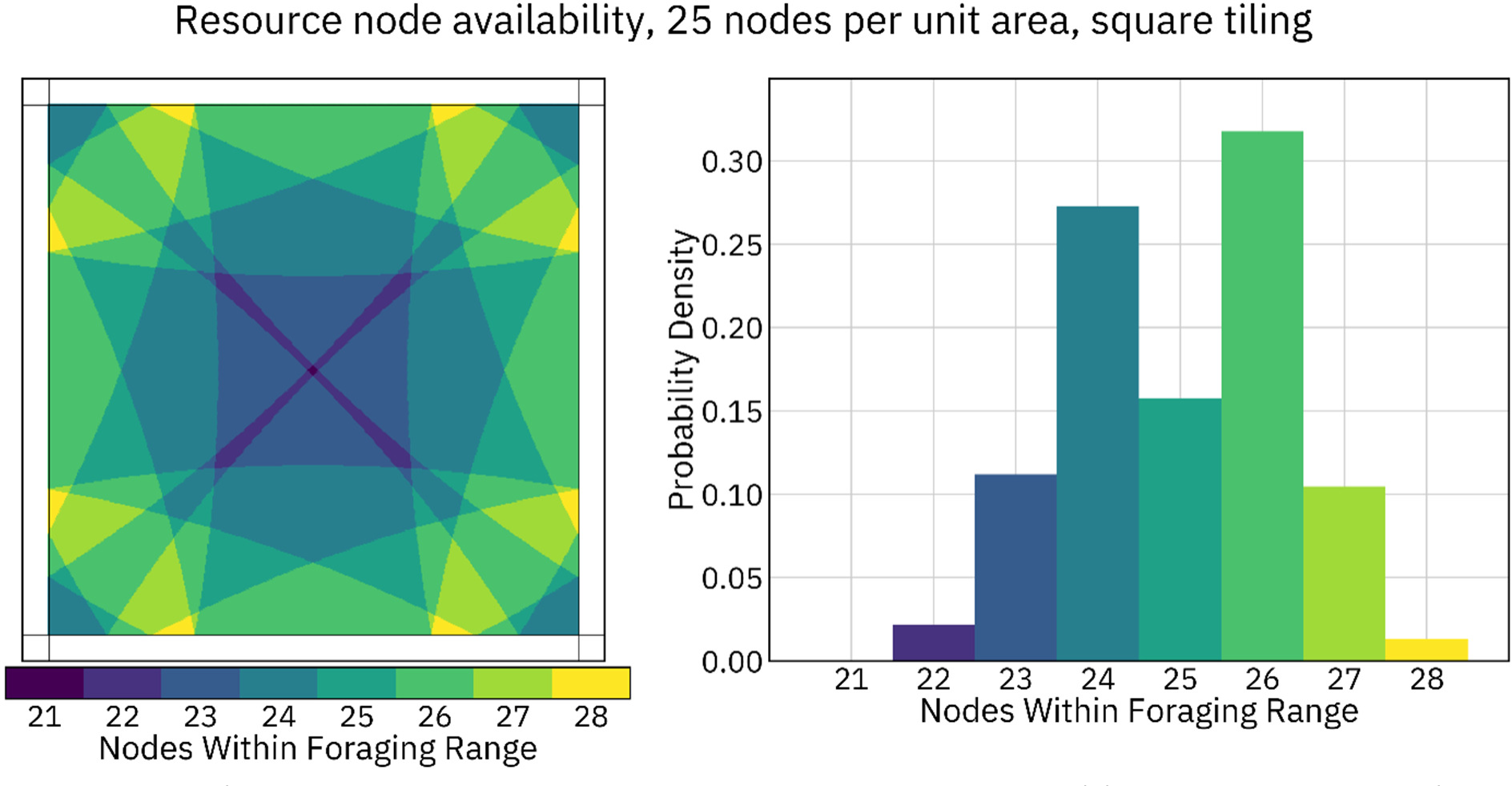
Unequal access to resource nodes, square tiling, 25 nodes per unit area. The number of resource nodes within the foraging radius of an individual depends on their location within a square tile. Individuals located in darker regions of the square (left panel) forage from fewer nodes, while those in brighter regions forage from more. The relative area of each color is depicted in the right panel. The mean value of this distribution is 25.00, and the standard deviation is 1.33 (equivalent to 5.3 percent of the foraging area).

**Supplemental Figure 16.**
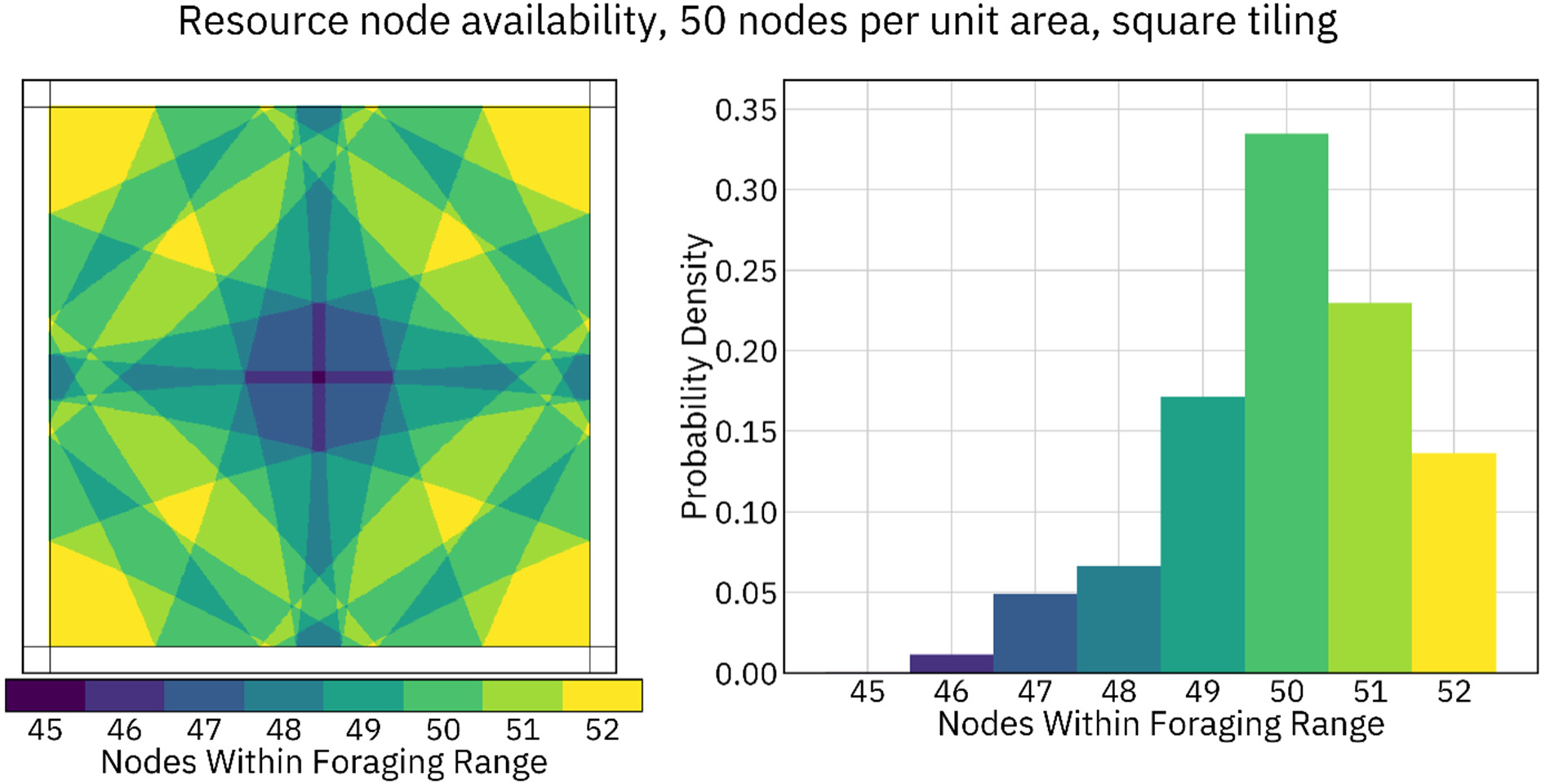
Unequal access to resource nodes, square tiling, 50 nodes per unit area. The number of resource nodes within the foraging radius of an individual depends on their location within a square tile. Individuals located in darker regions of the square (left panel) forage from fewer nodes, while those in brighter regions forage from more. The relative area of each color is depicted in the right panel. The mean value of this distribution is 50.00, and the standard deviation is 1.36 (equivalent to 2.7 percent of the foraging area).

**Supplemental Figure 17.**
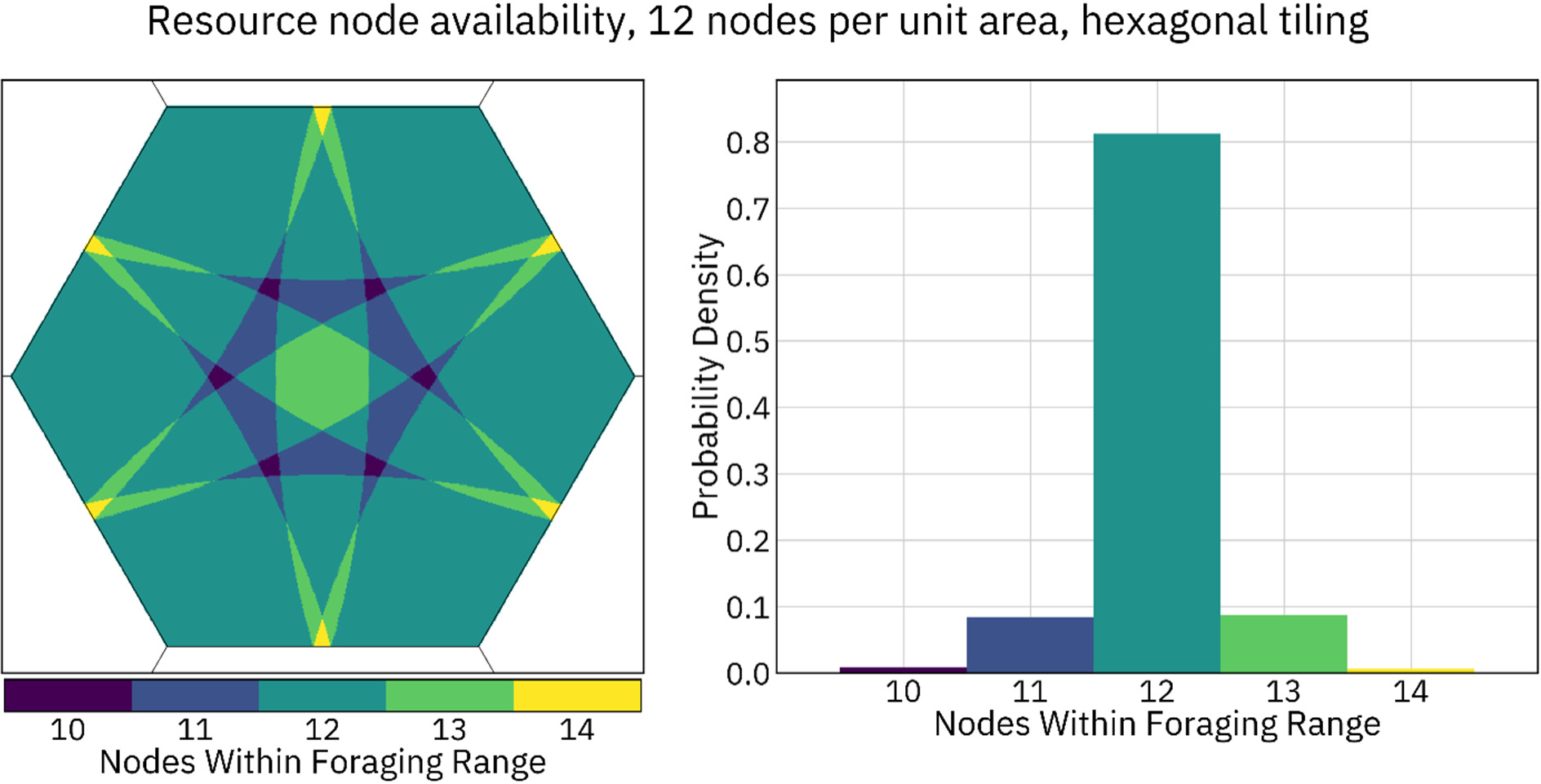
Unequal access to resource nodes, hexagonal tiling, 12 nodes per unit area. The number of resource nodes within the foraging radius of an individual depends on their location within a hexagonal tile. Individuals located in darker regions of the hexagon (left panel) forage from fewer nodes, while those in brighter regions forager from more. The relative area of each color is depicted in the right panel. The mean value of this distribution is 12.00, and the standard deviation is 0.48 (equivalent to a 4.0 percent of the foraging area).

**Supplemental Figure 18.**
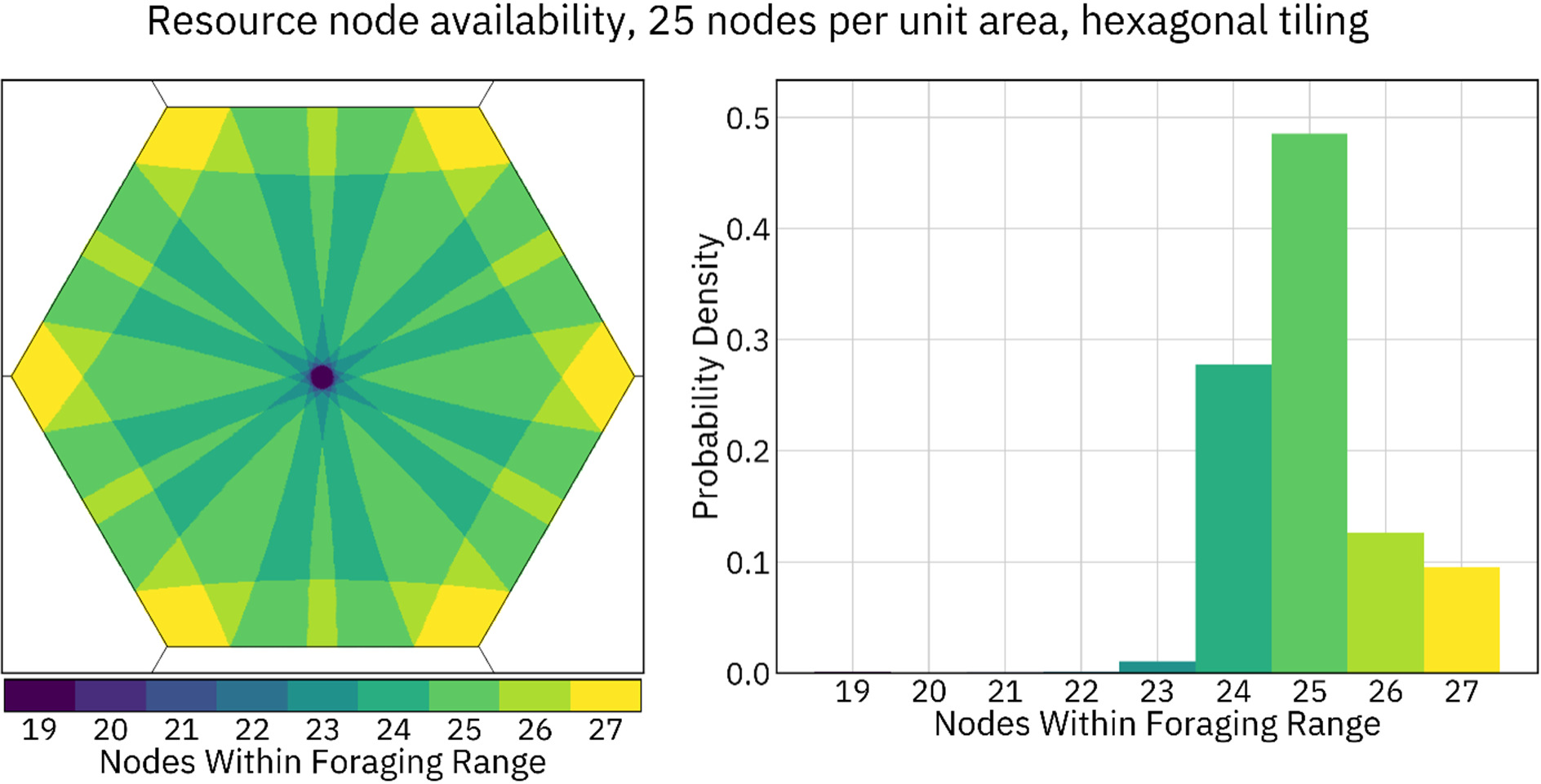
Unequal access to resource nodes, hexagonal tiling, 25 nodes per unit area. The number of resource nodes within the foraging radius of an individual depends on their location within a hexagonal tile. Individuals located in darker regions of the hexagon (left panel) forage from fewer nodes, while those in brighter regions forager from more. The relative area of each color is depicted in the right panel. The mean value of this distribution is 25.00, and the standard deviation is 0.96 (equivalent to a 3.8 percent of the foraging area).

**Supplemental Figure 19.**
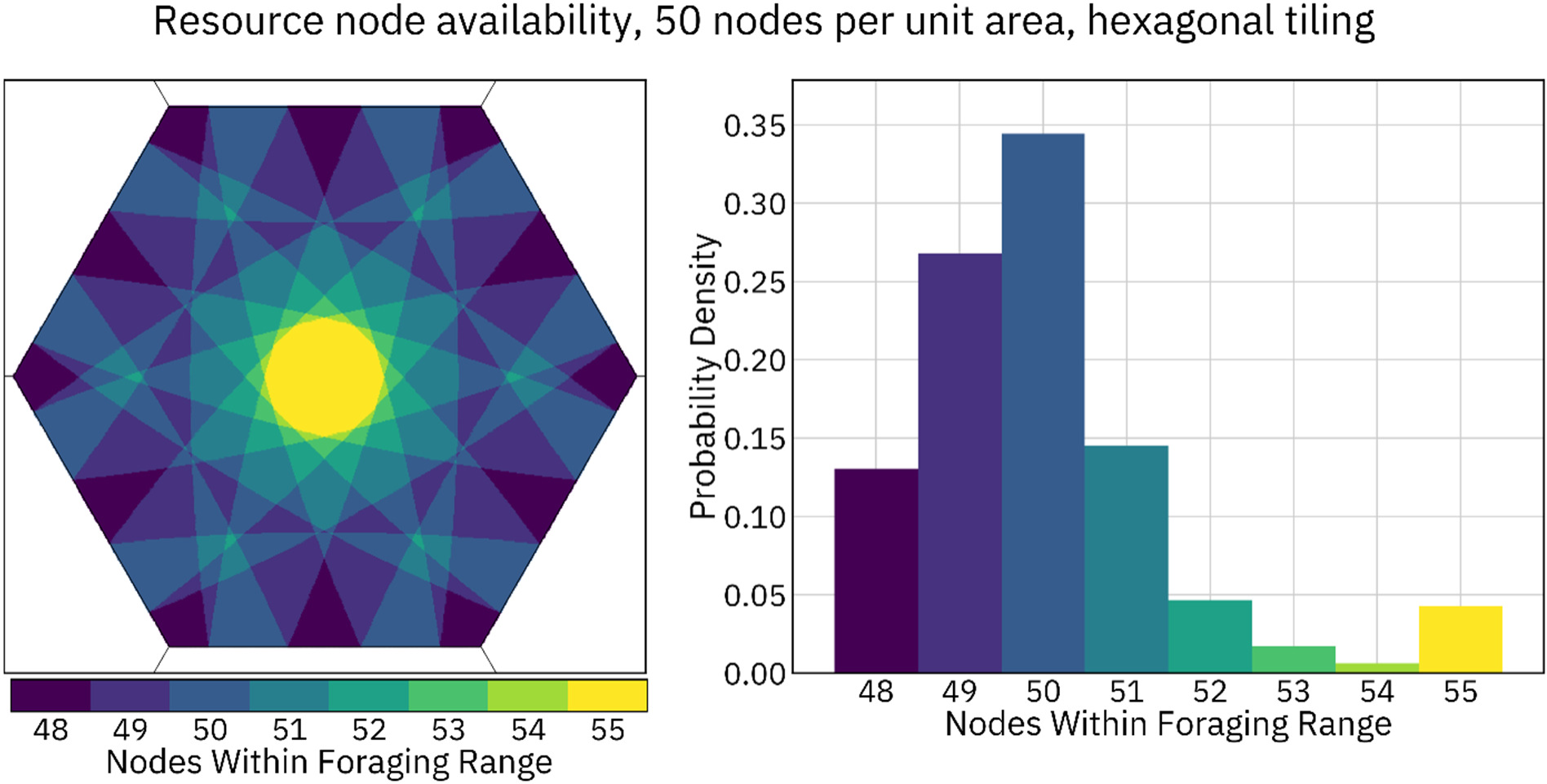
Unequal access to resource nodes, hexagonal tiling, 50 nodes per unit area. The number of resource nodes within the foraging radius of an individual depends on their location within a hexagonal tile. Individuals located in darker regions of the hexagon (left panel) forage from fewer nodes, while those in brighter regions forager from more. The relative area of each color is depicted in the right panel. The mean value of this distribution is 50.00, and the standard deviation is 1.56 (equivalent to a 3.1 percent of the foraging area).

**Supplemental Figure 20.**
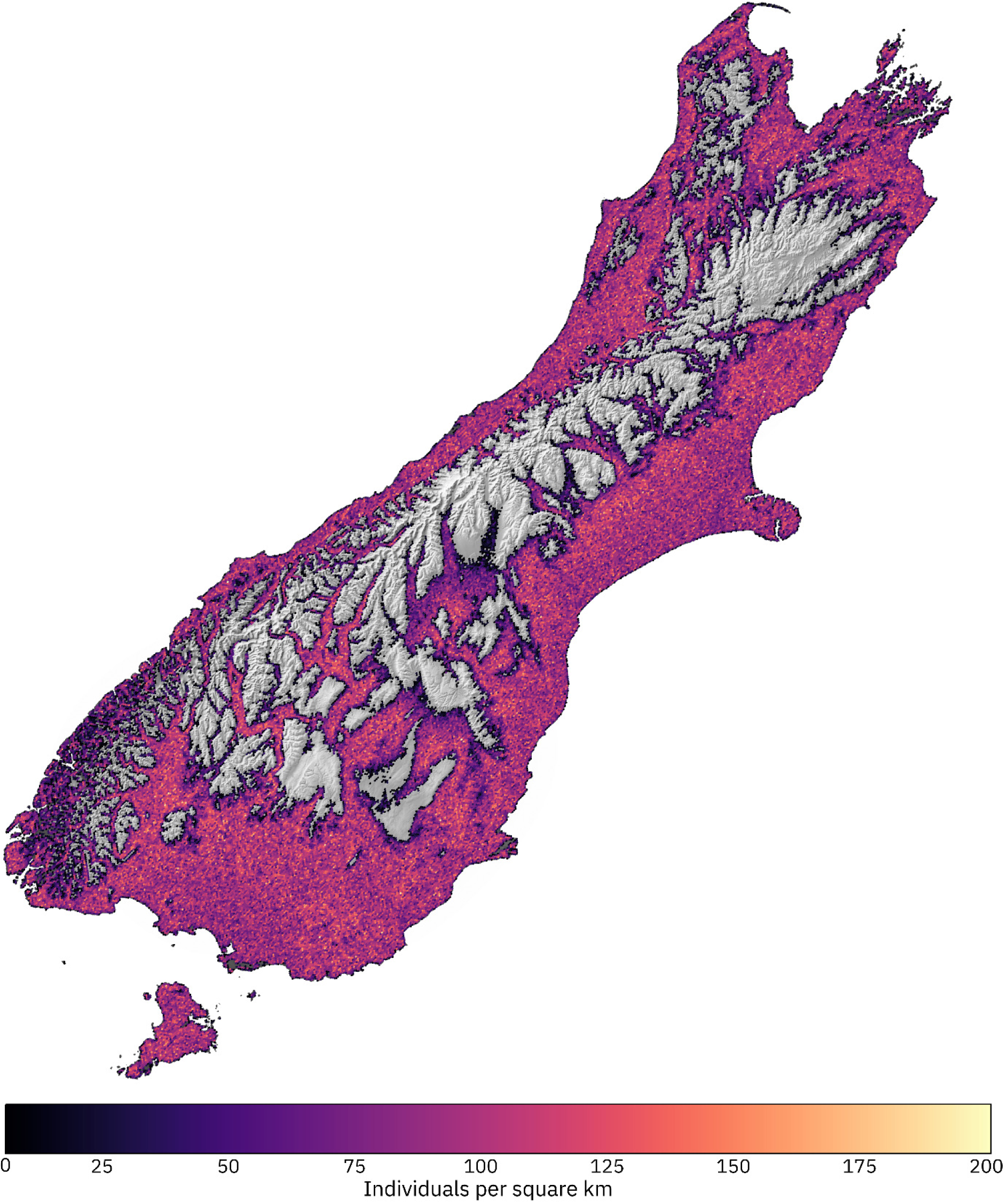
A heterogeneous landscape model of New Zealand. The modeled area was populated with a square grid of 4.8 million resource nodes and 10 million individuals. Habitat quality was defined as loosely inversely proportional to altitude, with the optimum habitat at about 300 meters above sea level. The color shade gradient denotes the density of individuals. Topographical map image (shown in black and white in areas with no resource nodes) courtesy NASA JPL.

**Supplemental Table 1.** Model runtimes, hexagonal tiling. Forty measurements of elapsed runtime per tick were collected and averaged for each method at each density and population size and at each of three node-placement densities. For the resource-explicit models depicted in this table, a hexagonal tiling of nodes was used. Color bars show the runtime of each model, on a different scale within each column, relative to the slowest runtime within that column.

| Model | Node density | Low density (20 individuals per unit area) |  | High density (200 individuals per unit area) |  |
| --- | --- | --- | --- | --- | --- |
|  |  | 100k individuals<br>Runtime per tick (s) | 1m individuals<br>Runtime per tick (s) | 100k individuals<br>Runtime per tick (s) | 1m individuals<br>Runtime per tick (s) |
| Panmictic | NA | 0.16 | 1.59 | 0.16 | 1.59 |
| Fixed-strength | NA | 3.03 | 38.98 | 17.01 | 233.43 |
| Linear | NA | 3.58 | 45.06 | 23.48 | 302.95 |
| Gaussian | NA | 5.76 | 67.88 | 44.81 | 523.23 |
| Inelastic | 12 | 0.83 | 9.42 | 1.18 | 15.71 |
|  | 25 | 0.98 | 11.31 | 1.33 | 17.53 |
|  | 50 | 1.33 | 15.05 | 1.57 | 21.27 |
| Inelastic "fair" | 12 | 1.00 | 11.77 | 1.35 | 18.16 |
|  | 25 | 1.33 | 15.81 | 1.66 | 22.75 |
|  | 50 | 2.04 | 23.50 | 2.26 | 31.15 |
| Inelastic<br>"resource-explicit<br>reproduction" | 12 | 0.98 | 10.95 | 0.82 | 10.59 |
|  | 25 | 1.38 | 15.71 | 1.18 | 15.20 |
|  | 50 | 2.14 | 24.40 | 1.91 | 24.10 |
| Elastic | 12 | 2.42 | 27.97 | 2.66 | 30.77 |
|  | 25 | 3.51 | 40.95 | 3.63 | 41.42 |
|  | 50 | 5.85 | 66.10 | 5.59 | 65.45 |

**Supplemental Table 2.** Model runtimes, random node placement. Forty measurements of elapsed runtime per tick were collected and averaged for each method at each density and population size and at each of three node-placement densities. For the resource-explicit models depicted in this table, nodes were randomly distributed across the landscape during each tick of the model. Color bars show the runtime of each model, on a different scale within each column, relative to the slowest runtime within that column.

| Model | Node density | Low density (20 individuals per unit area) |  | High density (200 individuals per unit area) |  |
| --- | --- | --- | --- | --- | --- |
|  |  | 100k individuals<br>Runtime per tick (s) | 1m individuals<br>Runtime per tick (s) | 100k individuals<br>Runtime per tick (s) | 1m individuals<br>Runtime per tick (s) |
| Panmictic | NA | 0.16 | 1.59 | 0.16 | 1.59 |
| Fixed-strength | NA | 3.03 | 38.98 | 17.01 | 233.43 |
| Linear | NA | 3.58 | 45.06 | 23.48 | 302.95 |
| Gaussian | NA | 5.76 | 67.88 | 44.81 | 523.23 |
| Inelastic | 12 | 0.95 | 11.94 | 1.27 | 16.77 |
|  | 25 | 1.29 | 17.10 | 1.50 | 20.32 |
|  | 50 | 1.97 | 27.03 | 1.92 | 26.57 |
| Inelastic "fair" | 12 | 1.23 | 15.44 | 1.51 | 19.65 |
|  | 25 | 1.91 | 24.05 | 2.00 | 26.11 |
|  | 50 | 3.23 | 41.09 | 2.98 | 38.72 |
| Inelastic<br>"resource-explicit<br>reproduction" | 12 | 1.19 | 14.67 | 0.92 | 12.40 |
|  | 25 | 1.88 | 24.37 | 1.38 | 19.27 |
|  | 50 | 3.15 | 44.24 | 2.41 | 32.62 |
| Elastic | 12 | 2.47 | 27.73 | 2.70 | 30.73 |
|  | 25 | 3.52 | 41.58 | 3.63 | 42.08 |
|  | 50 | 5.92 | 67.22 | 5.56 | 65.61 |

## REFERENCES

1. Wright, S. Evolution in Mendelian Populations. Genetics 16, 97 (1931).

2. Kingman, J. F. C. The coalescent. Stoch Process Their Appl 13, 235–248 (1982).

3. Pujolar, J. M. Conclusive evidence for panmixia in the American eel. Mol Ecol 22, 1761–1762 (2013).

4. Beveridge, M. & Simmons, L. W. Panmixia: an example from Dawson’s burrowing bee (*Amegilla dawsoni*) (Hymenoptera: Anthophorini). Mol Ecol 15, 951–957 (2006).

5. Brelsfoard, C. L. & Dobson, S. L. Population genetic structure of *Aedes polynesiensis* in the Society Islands of French Polynesia: Implications for control using a Wolbachia-based autocidal strategy. Parasit Vectors 5, 1–12 (2012).

6. Champer, J., Kim, I. K., Champer, S. E., Clark, A. G. & Messer, P. W. Suppression gene drive in continuous space can result in unstable persistence of both drive and wild-type alleles. Mol Ecol 30, 1086–1101 (2021).

7. Battey, C. J., Ralph, P. L. & Kern, A. D. Space is the Place: Effects of Continuous Spatial Structure on Analysis of Population Genetic Data. Genetics 215, 193 (2020).

8. Muktupavela, R. A. et al. Modeling the spatiotemporal spread of beneficial alleles using ancient genomes. Elife 11, (2022).

9. Champer, S. E., Kim, I. K., Clark, A. G., Messer, P. W. & Champer, J. *Anopheles* homing suppression drive candidates exhibit unexpected performance differences in simulations with spatial structure. Elife 11, (2022).

10. Lundgren, E. & Ralph, P. L. Are populations like a circuit? Comparing isolation by resistance to a new coalescent-based method. Mol Ecol Resour 19, 1388–1406 (2019).

11. Epperson, B. K. Geographical genetics. 356 (2003).

12. Birch, C. P. D., Oom, S. P. & Beecham, J. A. Rectangular and hexagonal grids used for observation, experiment and simulation in ecology. Ecol Modell 206, 347–359 (2007).

13. Garud, N. R., Messer, P. W., Buzbas, E. O. & Petrov, D. A. Recent Selective Sweeps in North American *Drosophila melanogaster* Show Signatures of Soft Sweeps. PLoS Genet 11, e1005004 (2015).

14. Min, J., Gupta, M., Desai, M. M. & Weissman, D. B. Spatial structure alters the site frequency spectrum produced by hitchhiking. Genetics 222, (2022).

15. Messer, P. W. & Petrov, D. A. Population genomics of rapid adaptation by soft selective sweeps. Trends Ecol Evol 28, 659–669 (2013).

16. Urban, M. C., Phillips, B. L., Skelly, D. K. & Shine, R. A toad more traveled: The heterogeneous invasion dynamics of cane toads in Australia. American Naturalist 171, 134–148 (2008).

17. Fisher, R. A. The Wave of Advance of Advantageous Genes. Ann Eugen 7, 355– 369 (1937).

18. Bentley, J. L. Multidimensional binary search trees used for associative searching. Commun ACM 18, 509–517 (1975).

19. Haller, B. C. & Messer, P. W. SLiM 3: Forward Genetic Simulations Beyond the Wright–Fisher Model. Mol Biol Evol 36, 632–637 (2019).

20. Haller, B. C. & Messer, P. W. SLiM 4: Multispecies eco-evolutionary modeling. Am Nat (2023) doi:10.1086/723601.

21. Byers, K. A., Lee, M. J., Patrick, D. M. & Himsworth, C. G. Rats About Town: A Systematic Review of Rat Movement in Urban Ecosystems. Front Ecol Evol 7, 13 (2019).

22. Haller, B.C., and Messer, P. W. SLiM: An Evolutionary Simulation Framework. http://benhaller.com/slim/SLiM_Manual.pdf (2016).

23. Eckhoff, P. A. A malaria transmission-directed model of mosquito life cycle and ecology. Malar J 10, 1–17 (2011).

24. Oberhauser, K. Suzanne Solensky, M. J., Journey North. & Monarch Larva Monitoring Project. Monarch butterfly biology & conservation. (Cornell University Press, 2004).

25. King, C. M. The Relationships between Beech (*Nothofagus* Sp.) Seedfall and Populations of Mice (*Mus musculus*), and the Demographic and Dietary Responses of Stoats (*Mustela erminea*), in Three New Zealand Forests. J Anim Ecol 52, 141 (1983).

26. Mantyka-Pringle, C. S., Martin, T. G. & Rhodes, J. R. Interactions between climate and habitat loss effects on biodiversity: a systematic review and meta-analysis. Glob Chang Biol 18, 1239–1252 (2012).

27. Pearse, I. S., Koenig, W. D. & Kelly, D. Mechanisms of mast seeding: resources, weather, cues, and selection. New Phytologist 212, 546–562 (2016).

28. Shopska, V., Denkova-Kostova, R. & Kostov, G. Modeling in Brewing—A Review. Processes 2022, Vol. 10, Page 267 10, 267 (2022).

29. Jones, C. G., Lawton, J. H. & Shachak, M. Organisms as Ecosystem Engineers. Oikos 69, 373 (1994).

30. Leer, B. van & Powell, K. G. Introduction to Computational Fluid Dynamics. Encyclopedia of Aerospace Engineering (2010) doi:10.1002/9780470686652.EAE048.

31. Barnes, J. & Hut, P. A hierarchical *O(N log N)* force-calculation algorithm. Nature 324, 446–449 (1986).

32. Grünbaum, B. & Shephard, G. C. Tilings by Regular Polygons. Mathematics Magazine 50, 227 (1977).

33. Champer, S. E. et al. Modeling CRISPR gene drives for suppression of invasive rodents using a supervised machine learning framework. PLoS Comput Biol 17, e1009660 (2021).

34. Statistics New Zealand. New Zealand Official Yearbook 2010. (David Bateman Ltd, 2010).

